# AI-assisted decoding of molecular signatures essential for NON-PHOTOTROPIC HYPOCOTYL3 condensation and function in phototropism

**DOI:** 10.1101/2024.06.17.598829

**Authors:** Prabha Manishankar, Lea Reuter, Leander Rohr, Atiara Fernandez, Yeliz Idil Yigit, Tanja Schmidt, Irina Droste-Borel, Jutta Keicher, Andrea Bock, Claudia Oecking

## Abstract

The plasma membrane-associated protein NON-PHOTOTROPIC HYPOCOTYL 3 (NPH3) is a key component of plant phototropism. Blue light induces the release of NPH3 into the cytosol, where it dynamically transitions into membrane-less biomolecular condensates. Here, we combine experimental data with the power of AI-based protein structure prediction to uncover a bipartite C-terminal motif that enables self-interaction of NPH3. We further demonstrate its importance for both the association of NPH3 with the plasma membrane and the assembly of condensates in the cytosol, with a different part of the bipartite motif playing the key role in each case. However, the formation of cytosolic condensates additionally requires the co-operative action of an N-terminal NPH3 signature. We propose that NPH3 forms a crosslinked 3D network in the cytosol based on distinct molecular determinants that simultaneously self-associate. NPH3 variants incapable of condensation are non-functional, suggesting a fundamental role of NPH3 phase separation for phototropism. This structural snapshot could have direct implications for future analyses of the plant-specific NPH3/RPT2-Like protein family.

## Introduction

The phototropic response describes the capability of plants to re-orient shoot growth towards a directional blue light (BL) source which ultimately enhances photosynthetic competence. The BL-activated and plasma membrane (PM)-associated AGCVIII kinases, phototropin1 (phot1) and phot2, function as primary photoreceptors. Asymmetric growth of the hypocotyl is based on the establishment of a lateral auxin gradient with a higher hormone concentration on the shaded side (Christie & Murphy, 2013; Fankhauser & Christie, 2015; Haga & Sakai, 2023; Küpers *et al*, 2020; Legris & Boccaccini, 2020). Accordingly, certain members of the PIN-FORMED (PIN) family of auxin efflux carriers are required for phototropic hypocotyl bending (Willige *et al*, 2013). In addition to the regulation of PIN activity by a family of AGCVIII protein kinases (D6 PROTEIN KINASES) (Willige *et al*, 2013; Zourelidou *et al*, 2014), alterations in the subcellular polarity of PIN proteins, particularly PIN3 (Ding *et al*, 2011; Zhang *et al*, 2017), are known to mediate the effect of BL on auxin distribution (Fankhauser & Christie, 2015; Hohm *et al*, 2013; Legris & Boccaccini, 2020). Although the mechanism linking photoreceptor activation to PIN protein modulation is still unknown, recent data have revealed that dynamic changes in the subcellular localization of the early phototropic signaling component NON-PHOTOTROPIC HYPOCOTYL 3 (NPH3) are crucial for phot1 signaling (Reuter *et al*, 2021; Sullivan *et al*, 2021). In darkness, NPH3 associates to the PM (Haga *et al*, 2015; Sullivan *et al*, 2019) in a phospholipid-dependent manner (Reuter *et al*, 2021) and physically interacts with the photoreceptor phot1 (Motchoulski & Liscum, 1999; Haga *et al*, 2015). Following BL perception, phot1 directly phosphorylates the third last residue (S744) of NPH3 (Sullivan *et al*, 2021), which represents a C-terminal 14-3-3 binding motif. Subsequent association of 14-3-3 proteins is causal for the light-induced release of NPH3 from the PM into the cytosol. This BL-induced relocation of NPH3 precedes its dynamic transitions into membrane-less condensate-like particles in the cytosol (Reuter *et al*, 2021; Sullivan *et al*, 2021). Importantly, the light-induced processes are reversible, eventually resulting in PM re-association of NPH3 (Haga *et al*, 2015; Reuter *et al*, 2021). Moreover, NPH3 mutants constitutively localizing to either the PM (incapable of 14-3-3 binding, e.g. NPH3_S744A) or condensates (impaired in PM association, e.g. NPH31′C51, NPH3_4K/A) show compromised functionality, consistent with a model where phototropic curvature critically depends on the 14-3-3 mediated dynamic change in NPH3 localization, ultimately leading to a gradient of NPH3 subcellular localization across the hypocotyl (Reuter *et al*, 2021; Sullivan *et al*, 2021).

Together with ROOT PHOTOTROPISM 2 (RPT2), NPH3 is the founding member of the plant-specific NPH3/RPT2-Like (NRL) protein family consisting of 33 members in Arabidopsis (Christie *et al*, 2018; Pedmale *et al*, 2010). Several of them act – comparable to NPH3 – as signal transducers in processes involving auxin (re)distribution in response to developmental or environmental cues, most likely via regulation of PIN auxin carriers (Cheng *et al*, 2007; Furutani *et al*, 2007; Li *et al*, 2011; Treml *et al*, 2005). Indeed, MAB4/ENP-Like (MEL) polypeptides were recently shown to maintain PIN polarity by limiting lateral diffusion (Glanc *et al*, 2021). Considering that several NRL proteins are potential direct substrate targets of the phot AGCVIII kinases (Sullivan *et al*, 2021; Waksman *et al*, 2023) and can interact with 14-3-3 via the corresponding phosphorylated motif (Reuter *et al*, 2021), phosphorylation-dependent 14-3-3 association may represent a more general regulatory mechanism. Noteworthy, MEL proteins also form particle-like structures in the cytosol (Furutani *et al*, 2007; Furutani *et al*, 2011; Glanc *et al*, 2021) that are reminiscent of NPH3 condensates, in addition to localization at the PM.

Membrane-less biomolecular condensates emerge as a fundamental mechanism to compartmentalize cellular functions. In plants, condensate assembly appears to link environmental cues to development (Emenecker *et al*, 2021; Field *et al*, 2023) and, as usual, is driven by multivalent macromolecular interactions (Banani *et al*, 2017; Lyon *et al*, 2021; Shin & Brangwynne, 2017). NRL family members are mostly characterized by a central NPH3 domain and two putative protein interaction domains, namely an N-terminal bric-a-brac, tramtrack and broad complex (BTB) domain and a C-terminal coiled-coil (CC) domain (Christie *et al*, 2018). To date, our understanding of the structure-function relationship is very limited, making it impossible to address the question of whether condensate formation of PM-detached NPH3 is essential for its action. Accurately modeling the structures of proteins and their complexes using artificial intelligence (AI) is revolutionizing molecular biology (Evans *et al*, 2022; Jumper *et al*, 2021). Here, we closely link experimental data and AI-based protein structure predictions to decipher the molecular functions of the structured NPH3 domains. We report the identification of a bipartite C-terminal self-association motif that enables formation of homo-oligomers, likely trimers, and is required for both PM association of NPH3 in darkness and BL-triggered condensate assembly in the cytosol. The bipartite motif consists of the CC domain and a yet undescribed Linear Interacting Peptide (LIP) at the C-terminal end of the NPH3 domain. Remarkably, only the LIP-mediated self-association of NPH3 with low affinity is essential for the transition to the condensed state. In addition, the N-terminal BTB domain contributes directly to the oligomeric states of NPH3 in cytoplasmic assemblies.

Overall, we propose the conversion of a strict C-terminal self-association of NPH3 at the PM into a higher-order 3D self-crosslinking in the cytosol mediated by co-operative self-association of both the LIP motif and the BTB domain. Several NPH3 variants incapable of condensate formation are non-functional, suggesting that the temporal regulation of the BL-induced NPH3 cycling between the PM and the cytosol is critical for its function in auxin-dependent phototropism.

## Results

### NPH3 self-associates independently of its subcellular localization and its BTB domain

According to protein signature models NPH3 possesses an N-terminal BTB, a central NPH3 and a C-terminal CC domain (Christie *et al*, 2018). Structure prediction by AI-based tools AlphaFold2 (AF2) (Jumper *et al*, 2021) or AF3 (Abramson *et al*, 2024), confirmed the above domains, each characterized by defined secondary structures and high confidence scores (per residue predicted Local Distance Difference Test, pLDDT). In addition, six intrinsically disordered regions (IDR), showing low pLDDT values and covering about 40% of the protein, are distributed along the entire length of NPH3 (Figs. 1A and EV1A). Nevertheless, the positions (Predicted Aligned Error, PAE) of the structured domains are well-defined relative to each other – with the exception of the CC domain (Fig. EV1B).

**Figure 1.**
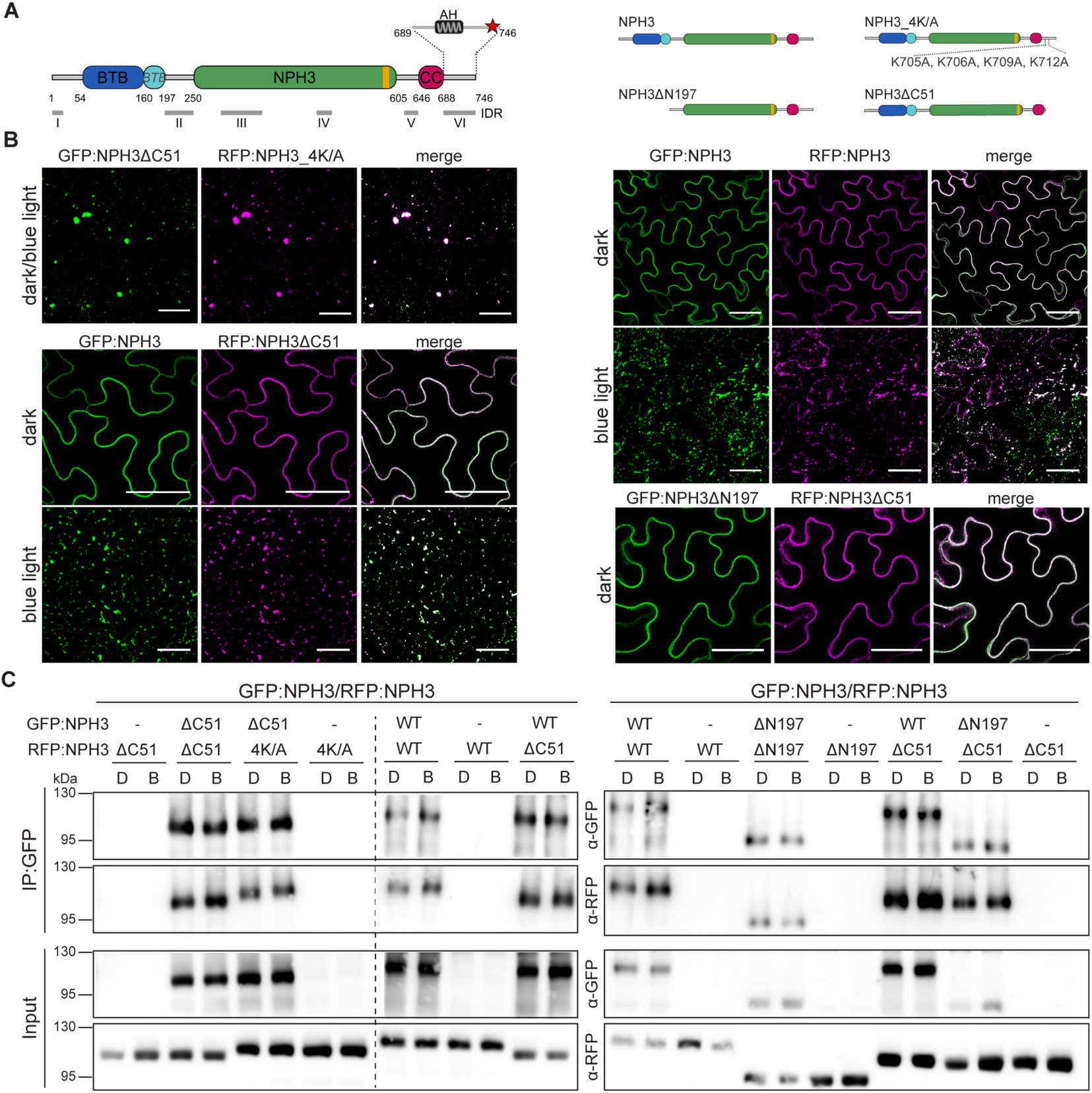
NPH3 self-associates at the PM and within cytosolic condensates independent of its BTB domain. (**A**) Domain architecture of NPH3 (746 amino acids) and location of IDRs (AF2 prediction). C-terminal BTB extension highlighted in light blue (*BTB*), LIP motif highlighted in yellow, AH amphipathic helix, see text for details **(left panel).** Schematic representation of NPH3 constructs used **(right panel)**. (**B**) Representative confocal images of leaf epidermal cells from *N. benthamiana* transiently co-expressing different GFP/RFP:NPH3 variants (35S promoter). Dark-adapted tobacco plants were either kept in darkness or treated with BL (∼ 11 min GFP laser). Z-stack projections of BL-treated samples are shown. Scale bars, 50 µm. These experiments were repeated at least three times with similar results. (**C**) *In vivo* interaction of GFP/RFP:NPH3 variants transiently co-expressed (35S promoter) in *N. benthamiana* leaves. Dark-adapted tobacco plants were either kept in darkness (D) or treated with BL (B) (10 µmol m^−2^s^−1^, 1h). The crude protein extract was immunoprecipitated using GFP beads. Immunodetection of input and immunoprecipitated proteins (IP:GFP) is shown. Experiments were performed at least two times with similar results. Dotted line indicates that the same blot has been sliced and the sides swapped.

Protein condensate formation relies on multivalent interactions. Since NPH3 variants that constitutively form cytosolic condensates (NPH3ΔC51, NPH3_4K/A) strictly co-localize when transiently co-expressed in *N. benthamiana* leaves (Reuter *et al*, 2021) (Fig. 1B), we wondered whether NPH3 associates with itself in cytoplasmic assemblies. Stringent co-immunoprecipitation (CoIP) of fluorophore-tagged proteins confirmed their physical association (Fig. 1C). However, NPH3 self-association is not restricted to cytosolic assemblies but also occurs at the PM (NPH3/NPH3 darkness) (Fig. 1 B,C). Remarkably, condensate-localizing RFP:NPH3ΔC51 uniformly associated with the PM in the dark when co-expressed with GFP:NPH3 (Fig. 1B), implying that self-association of NPH3 enables the relocation of cytosolic variants to the PM. This phenomenon is hereafter referred to as self-association linked to relocation. Subsequent BL irradiation triggered co-localization of RFP:NPH3ΔC51 and GFP:NPH3 in cytosolic assemblies (Fig. 1B). Conversely, PM retention in BL was observed when RFP:NPH3ΔC51 or RFP:NPH3 were co-expressed with the constitutively PM-localized GFP:NPH3_S744A mutant (Reuter *et al*, 2021; Sullivan *et al*, 2021) (Fig. EV1C). In each case, association of NPH3 variants was validated by CoIP (Figs. 1C and EV1D). The BTB domain has often been associated with homo- or hetero-multimerization of NRL family members (Christie & Murphy, 2013; Inada *et al*, 2004; Pedmale *et al*, 2010; Upadhyay-Tiwari *et al*, 2024). However, deletion of the BTB domain (NPH3ΔN197) had no effect on self-association at the PM or the ability to translocate RFP:NPH3ΔC51 to the PM in darkness (Figs. 1B,C and EV1C). Consequently, BTB-independent self-association of NPH3 at the PM (dark) might be coupled with the dynamic formation of higher-order homo- or hetero-oligomers in the cytosol (BL), ultimately leading to condensate formation.

### AlphaFold2-multimer assisted identification of a bipartite C-terminal self-association motif

We next examined whether the CC domain of NPH3 contributes directly to self-association linked to relocation. When NPH3 lacking the CC domain (NPH3ΔC104) was co-expressed with NPH3 in darkness, it formed condensate-like particles near the PM (Fig. 2A) – an obvious difference to the uniform PM localization of NPH3ΔC51 (Fig. 1B). In BL, however, NPH3ΔC104 behaves like NPH3ΔC51, as it co-localizes with NPH3 in cytosolic condensates. Co-localization in condensates was also observed when NPH3ΔC104 was co-expressed with NPH3ΔC51 or itself (Figs. 2A and EV2A). Semi-quantitative CoIP revealed that NPH3ΔC104 – when compared to NPH3ΔC51 - associated significantly weaker with NPH3, NPH3ΔC51 or itself, although complex formation was still detectable (Figs. 2B and EV2B). We thus hypothesized that the CC domain promotes a high-affinity interaction essential for self-association linked to relocation. By contrast, the remaining weak interaction capability of NPH3ΔC104 might be sufficient to contribute to multivalency-driven condensate assembly. To unveil potential homomeric NPH3 interfaces, AF2-Multimer (AF2M), a protein complex structure prediction tool (Evans *et al*, 2022), and AF3 (Abramson *et al*, 2024) were used. Given the experimental evidence for the importance of the CC (Fig. 2B and EV2B), particular attention was paid to the confidence score for this domain. On this basis, NPH3 will most likely form a trimer (Fig. EV2C), which serves as a lead structure for a homooligomer (≥3) in the following. Confidence scores (pLDDT) of the AF3 prediction were lower than those of AF2M – probably due to the multitude of IDRs in NPH3 (see https://alphafoldserver.com, limitations of the AF3 model). As the models do not differ in principle, the AF2M predictions are shown below. Trimer formation of NPH3 is facilitated by small interfaces, particularly the CC super helices (Figs. 2C and EV3A), which are likely to be located near the PM, as the amphipathic helix required for PM association (arrows Fig. 2C) is found in the adjacent region (Reuter *et al*, 2021). Excitingly, upstream of the CC trimer, AF2M predicts another three-helical bundle. Here, a Linear Interacting Peptide (LIP) (MobiDB: _592_LRV**VVQVLF**, highlighted in yellow in all ribbon plots and schemes of NPH3) might establish hydrophobic helix-helix interfaces. The so far unrecognized LIP represents the (almost) C-terminal end of the NPH3 domain, exhibiting high sequence identity within the NRL family (Motchoulski & Liscum, 1999) (Fig. EV3B). In each trimeric NPH3 subunit, the entire region encompassing the BTB and NPH3 domains, including LIP, adopts a conformation close to that of the NPH3 monomer, while the position of the CC appears to be flexible in relation to the rest of the protein (PAE plot, Fig. EV3A). Due to the putative LIP interactions, the NPH3 trimer exhibits an almost perfect rectangular arrangement of its monomeric subunits. A view from below shows a symmetrically arranged 3-bladed propeller with the LIP-trimer in its center (Fig. 2C bottom).

**Figure 2.**
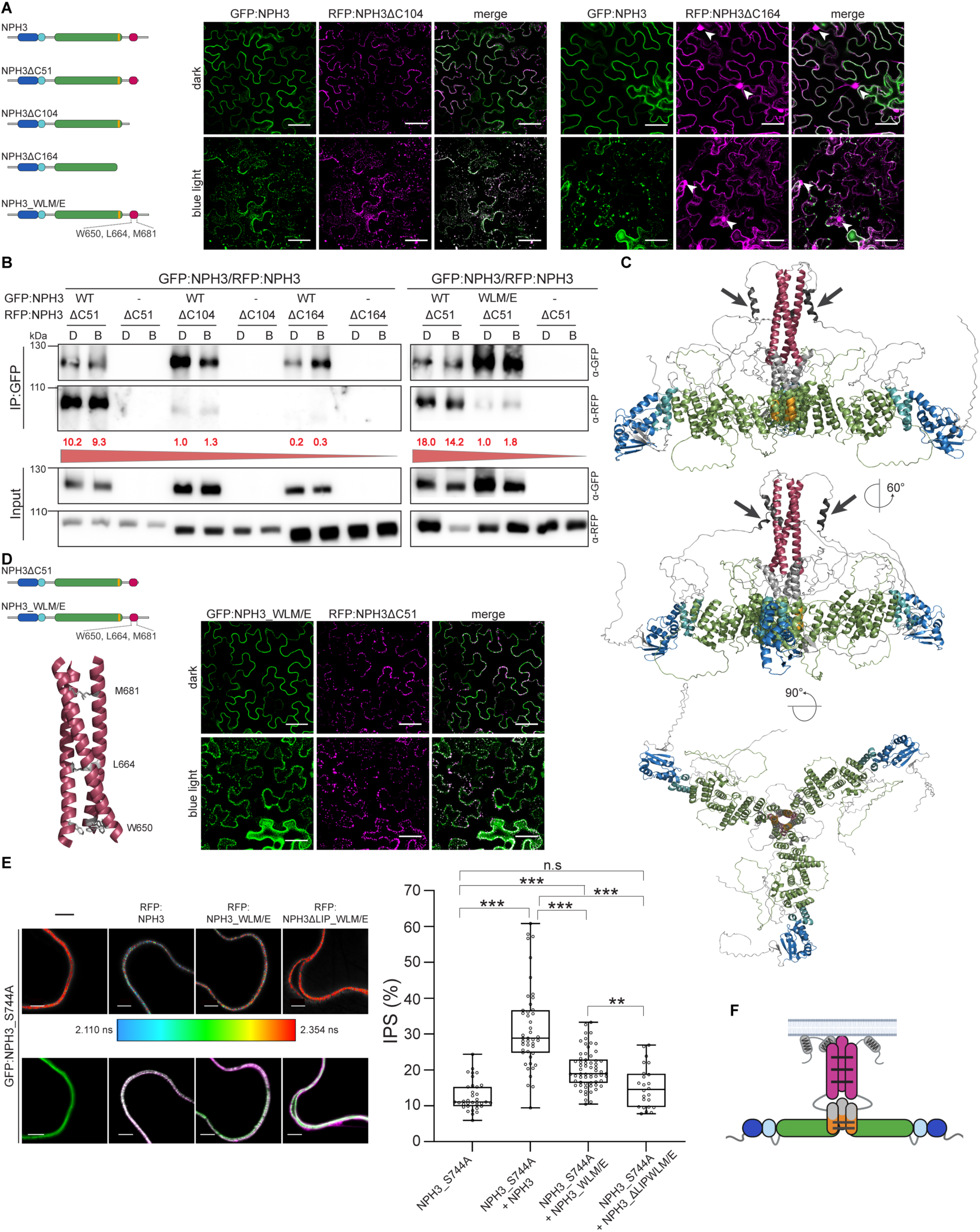
NPH3 is characterized by a bipartite C-terminal self-association motif. (**A**, **D**) Schematic representation of NPH3 variants used and representative confocal images of leaf epidermal cells from *N. benthamiana* transiently co-expressing (35S promoter) different GFP/RFP:NPH3 variants. Dark-adapted tobacco plants were either kept in darkness or treated with BL (∼ 11 min GFP laser). Z-stack projections of BL-treated samples are shown. Scale bars, 50 µm. Experiments were repeated at least three times with similar results. White arrowheads indicate nucleus. (**B**) *In vivo* interaction of GFP/RFP:NPH3 variants transiently co-expressed (35S promoter) in *N. benthamiana* leaves. Dark-adapted tobacco plants were either kept in darkness (D) or treated with BL (B) (10 µmol m^−2^s^−1^, 1h). The crude protein extract was immunoprecipitated using GFP beads. Immunodetection of input and immunoprecipitated proteins (IP:GFP) is shown. Experiments were performed at least two times with similar results. Values in red: normalized RFP/GFP intensity ratio. (**C**) Ribbon plot of the NPH3 trimer predicted by AF2M and color-coded according to the schematic domain representation (Fig. 1A). Grey arrow: amphipathic helix. (**E**) Interaction Pixels Analysis based on FRET-FLIM measurements performed on leaf epidermal cells from dark-adapted *N. benthamiana* transiently (co-)expressing (35S promoter) GFP:NPH3_S744A and RFP-tagged NPH3 variants. Representative color-coded FLT images (2.354 ns: mean FLT of donor-only, 2.110 ns: IPS threshold) (**upper panel**) and merged confocal images (**lower panel**) are shown. Scale bars, 5 µm. Statistical differences in IPS were analyzed using Kruskal-Wallis test, followed by a Dunn post hoc analysis with Holm-adjusted p-values (** p < 0.01, *** p < 0.001, n.s. non significant) (**right panel**). Center line: median, bound of boxes: minima and maxima (25^th^ and the 75^th^ percentiles), whiskers: 1.5 x IQR. Number of cells analyzed (from left to right): 35, 45, 63, 24. (**F**) Schematic model of NPH3 illustrating the self-association of NPH3 at the PM mediated by the bipartite C-terminal motif. Color-coding according to the schematic domain representation (Fig. 1A). Dark and light grey bars indicate high and low affinity, respectively.

We first validated CC-mediated homo-oligomerization of NPH3 by size exclusion chromatography of recombinant C-terminal NPH3 fragments tagged with (His)_6_-maltose-binding protein (MBP). While the MBP:NPH3-C51 (C-terminal 51 residues) fusion protein (MW: 48.2 kDa) eluted at a volume indicative of a monomer (15 ml, approx. 47 kDa), the presence of the CC domain (MBP:NPH3-C104, C-terminal 104 residues, MW: 54.4 kDa) resulted in a significant shift in the elution peak, suggesting a native molecular weight consistent with a trimer (12 ml, approx. 166 kDa), with a small fraction detectable as a hexamer (10.5 ml, approx. 346 kDa) (Fig. EV3C). To destabilize the CC superstructure, three hydrophobic residues (Fig. 2D: W650, L664, M681) were replaced with glutamic acid (NPH3_WLM/E). Indeed, MBP:NPH3-C104_WLM/E significantly impaired homo-oligomerization of the CC, likely resulting in a monomeric state (14 ml, approx. 74 kDa) (Fig. EV3C).

Based on this, we tested our hypothesis of CC-mediated high-affinity self-association of NPH3 linked to relocation. In contrast to GFP:NPH3, GFP:NPH3_WLM/E was unable to uniformly relocate RFP:NPH3ΔC51 to the PM in darkness (Fig. 2D), and its ability to bind NPH3ΔC51 was markedly reduced (Fig. 2B). However, upon BL treatment, GFP:NPH3_WLM/E and NPH3ΔC51 exhibited co-localization in cytoplasmic assemblies (Fig. 2D). Overall, this reflects the concurrent expression of PM-associated NPH3 and cytosolic NPH3ΔC104, which lacks the CC domain (Fig. 2A). We therefore concluded that the CC domain is essential for the high-affinity self-association of NPH3 linked to relocation. According to the structural prediction, the LIP motif may represent the molecular signature allowing for weak interaction and potentially condensate formation of NPH3 variants impaired in CC-mediated high-affinity interaction (NPH3ΔC104, NPH3_WLM/E). Notably, AF2M still predicts the formation of a trimer for NPH3ΔC104, with high confidence scores for LIP residues at the trimer interface (Fig. EV3D). By contrast, trimer formation is not supported by AF2M if the LIP motif is additionally truncated (NPH3ΔC164). Co-expression analyses demonstrated a preferential cytosolic localization of NPH3ΔC164 and a clear separation from the strictly condensate-forming NPH3ΔC51 as well as from NPH3 both in darkness and upon BL irradiation (Figs. 2A and EV2A). Moreover, semi-quantitative CoIP analyses revealed a clear stepwise decrease in avidity: compared to NPH3ΔC51, deletion of the CC (NPH3ΔC104) led to a significantly reduced but still detectable association, while additional truncation of the LIP motif (NPH3ΔC164) almost abolished complex formation with NPH3 or NPH3ΔC51. A comparable gradual decrease in interaction was also observed for the co-expression of the respective C-terminally deleted NPH3 variants (Figs. 2B and EV2A,B).

To substantiate the involvement of the LIP motif in self-association of NPH3, *in planta* Förster resonance energy transfer (FRET) by fluorescence lifetime imaging (FLIM) was performed. Given the mobility of condensates, constitutively PM-associated GFP:NPH3_S744A was used as donor, while RFP-tagged NPH3 variants served as acceptors. The percentage of interaction pixels (IPS), defined by strongly reduced fluorescence lifetime (FLT) of GFP (Bücherl *et al*, 2013), was quantified. A notable increase in IPS was observed when RFP-tagged NPH3 was co-expressed with the donor (32% IPS, compared to 13% IPS of donor-only). Destabilization of the CC-superstructure resulted in a reduced IPS proportion (RFP:NPH3_WLM/E, 21% IPS), although this was still significantly higher than the donor-only control. In contrast, no substantial difference in IPS was detected when, in addition to CC destabilization, self-association of the LIP motif was impaired (RFP:NPH3ΔLIP_WLM/E, 15% IPS) (Fig. 2E). In this mutant variant, deletion of the 9 aa LIP motif (_592_LRV**VVQVLF**) served to prevent low-affinity homo-oligomerization.

Taken together, these results suggest a bipartite C-terminal self-association motif in NPH3 (Fig. 2F), consisting of the CC domain, which mediates high-affinity homo-oligomerization required for relocation, and the LIP motif, which facilitates a low-affinity interaction that appears sufficient for transitioning NPH3 into a condensed state in the cytosol.

### The bipartite C-terminal self-association motif is required for plasma membrane recruitment of NPH3 in darkness

The region downstream of the CC, NPH3-C51, contains the amphipathic helix crucial for phospholipid-dependent PM association (Figs. 1A and 2C arrows). Despite its direct binding to phospholipids *in vitro* (Reuter *et al*, 2021), RFP:NPH3-C51 surprisingly failed to associate with the PM *in vivo* (Fig. 3A,B). However, extension by the CC domain (RFP:NPH3-C104) enabled PM association in darkness, as confirmed by subcellular fractionation (Fig. 3B,C). Membrane targeting of RFP:NPH3-C104 was in turn impaired when formation of CC homooligomers (RFP:NPH3-C104_WLM/E) or alternatively phospholipid binding of NPH3 (RFP:NPH3-C104_4K/A, see (Reuter *et al*, 2021)) was reduced (Fig. 3B,C). Overall, PM recruitment of NPH3-C104 in darkness requires both homo-oligomerization and phospholipid interaction.

**Figure 3.**
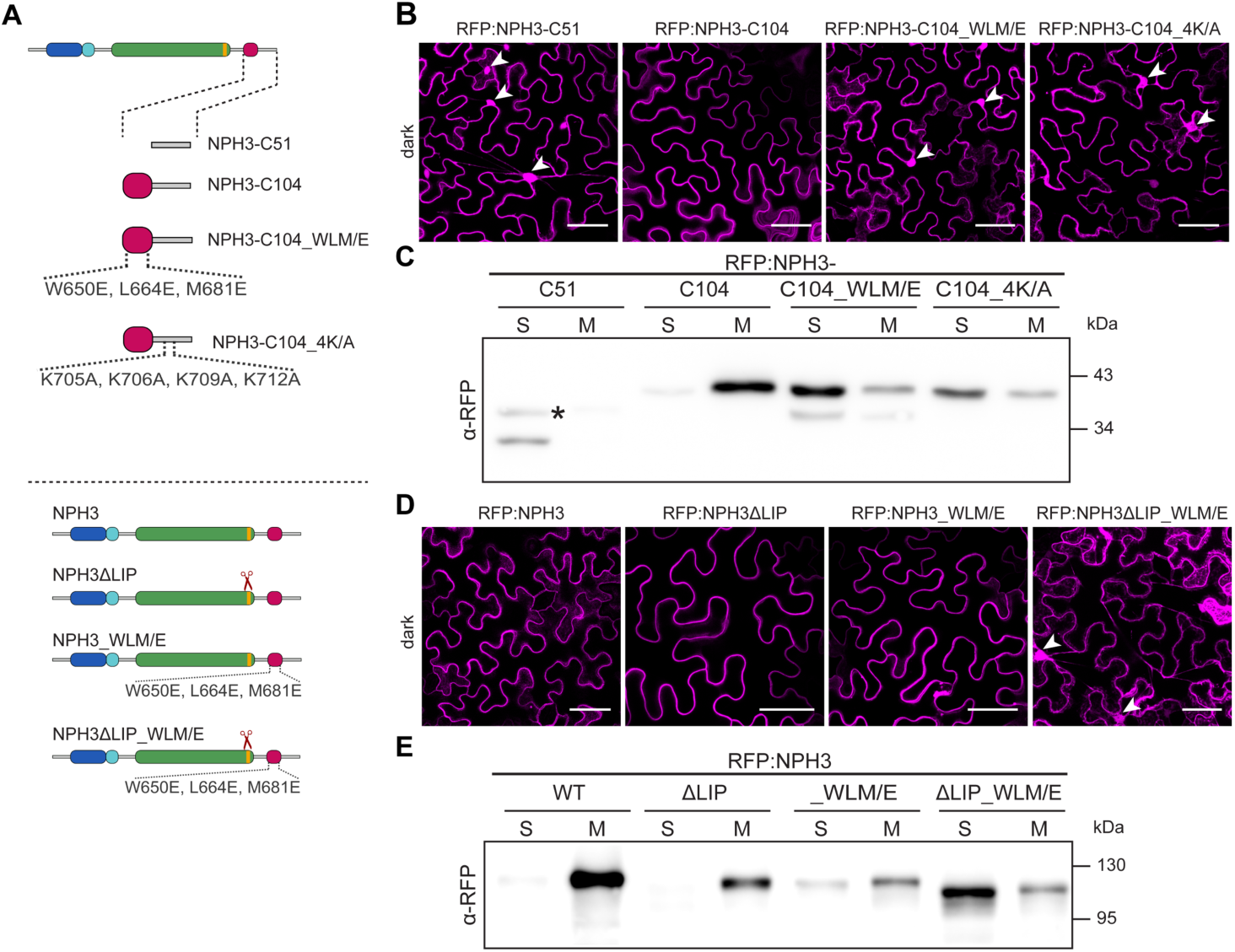
Homo-oligomerization mediated by the bipartite motif is essential for PM association of NPH3 *in vivo*. (**A**) Schematic representation of C-terminal NPH3 fragments and NPH3 variants used. (**B, D**) Representative confocal images of leaf epidermal cells from dark-adapted *N. benthamiana* transiently expressing (35S promoter) RFP:NPH3 variants. Z-stack projections are shown. Scale bars, 50 µm. White arrowheads indicate nucleus. (**C, E**) Representative α-RFP immunoblots following subcellular fractionation of protein extracts prepared from dark-adapted *N. benthamiana* leaves transiently expressing (35S promoter) RFP:NPH3 variants. Proteins in each fraction (∼ 10 µg) were separated on SDS-PAGE gels. It is noteworthy that the total amount of soluble proteins (S) is ∼10 times higher as compared to the total amount of microsomal proteins (M). (*) Asterisk indicates non-specific band. All experiments were performed at least three times with comparable results.

Analysis of full-length NPH3 variants aimed to understand the role of the LIP motif in PM attachment. Deletion of the 9 aa comprising LIP motif (_592_LRV**VVQVLF**) served as a means to prevent the LIP-mediated low-affinity homo-oligomerization (RFP:NPH3ΔLIP) and had no apparent effect on PM association (Fig. 3D,E). Disruption of the homotypic CC interaction shifted a proportion of RFP:NPH3_WLM/E to the soluble fraction following subcellular fractionation, but the shift was substantially smaller compared to NPH3-C104_WLM/E, which lacks the LIP motif (Fig. 3C,E). Remarkably, the full-length RFP:NPH3_WLM/E clearly localized to the cell periphery, suggesting weak PM attachment *in vivo,* likely due to LIP-mediated low-affinity self-association. In fact, a significant impairment of PM recruitment was only observed when the self-association of both parts of the bipartite structured motif was disturbed (RFP:NPH3ΔLIP_WLM/E) (Fig. 3D,E).

We therefore concluded that the bipartite C-terminal self-association motif acts unequally redundant for NPH3 membrane recruitment, with the CC domain fully compensating for loss of the LIP motif. However, when CC homo-oligomerization is impaired, the LIP motif partially compensates and facilitates weak PM association of NPH3, probably due to low-affinity self-association.

### Molecular NPH3 signatures essential for condensate formation

Biomolecular condensates typically form via liquid-liquid phase separation, and their formation relies on multivalent molecules. Multivalency in proteins can be achieved by structured interaction domains, IDRs or a combination of both (Banani *et al*, 2017). We have recently reported that BL irradiation initially triggers re-localization of NPH3 from the PM to the cytosol, where it dynamically transitions to a condensed state. While 14-3-3 association is critical for initial PM detachment of NPH3, it is not required for condensate assembly as shown by the analysis of various NPH3 mutant proteins, including full-length variants (Reuter *et al*, 2021). Phase separation may therefore be due to intrinsic properties of NPH3 when a critical concentration in the cytosol is exceeded.

Our co-expression data suggested that deletion of the C-terminal bipartite motif prevents condensate formation (NPH3ΔC164) (Fig. 2A). Single expression of NPH3ΔC164 confirmed the direct contribution of the bipartite motif to the oligomeric states of NPH3 in cytoplasmic assemblies (Figs. 4A,B). However, exclusive interference with CC formation (NPH3ΔC104, NPH3_WLM/E) only marginally affected condensate assembly of NPH3, and, unexpectedly, deletion of the LIP motif alone (NPH3ΔLIP) abolished transition of cytosolic NPH3 to the condensed state (Fig. 4A,B). Replacement of four hydrophobic/uncharged residues potentially involved in hydrophobic LIP-LIP interfaces with alanine or glutamic acid (_592_<u>L</u>RV**V<u>V</u>QV<u>L</u>F**SE<u>Q</u> ⇒ _592_<u>E</u>RV**V<u>A</u>QV<u>E</u>F**SE<u>A</u>) (NPH3_4LVQ/EA, Fig. EV4A) revealed that lack of homotypic LIP interactions is causal for the loss of phase separation ability of NPH3ΔLIP (Fig. 4A). Taken together, the CC domain and the LIP motif act non-redundantly in condensate formation of NPH3: here, the LIP motif is instrumental, while the CC is dispensable.

**Fig. 4.**
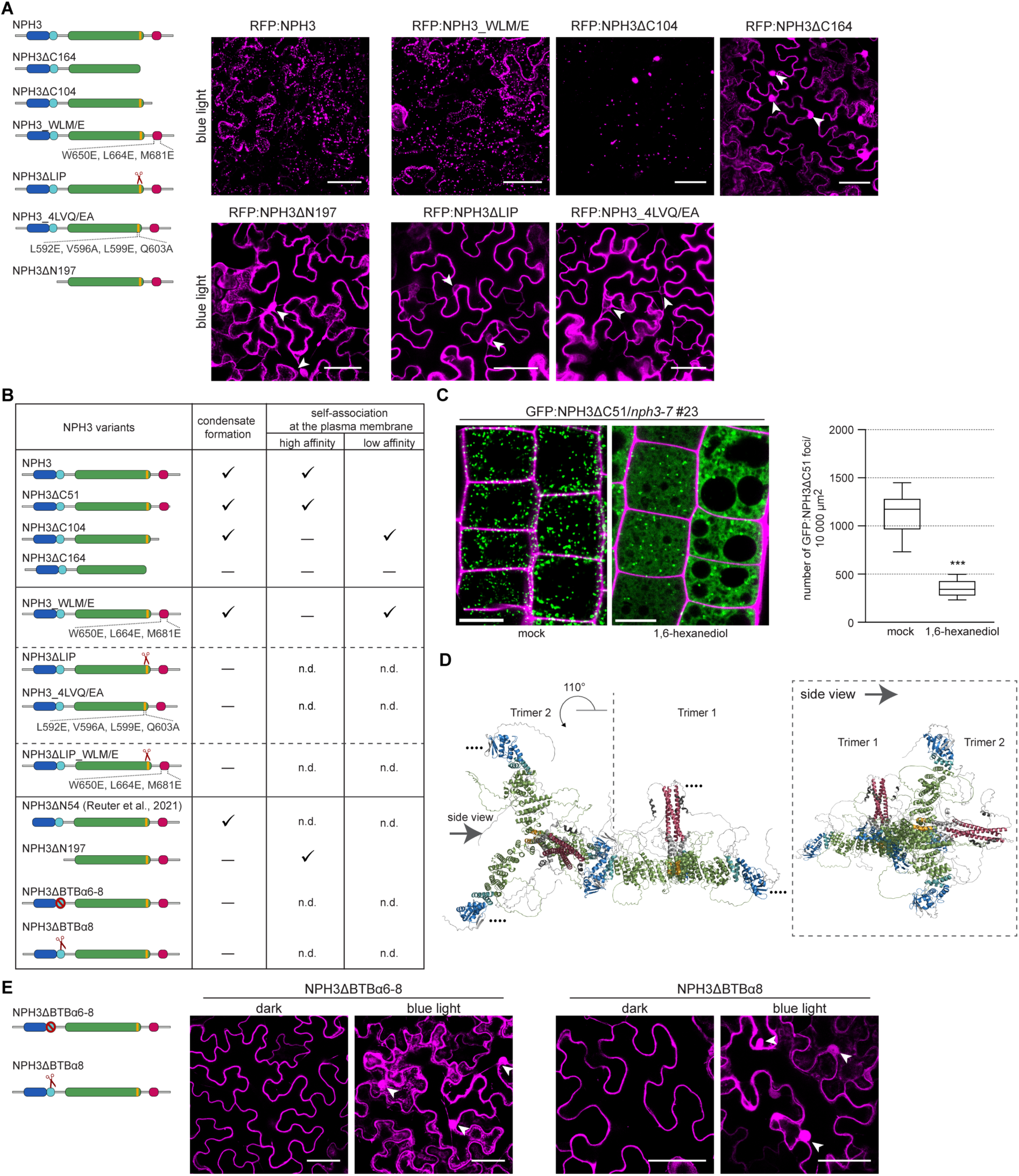
Condensate assembly of NPH3 requires the LIP motif and the BTB domain. (**A, E**) Schematic representation of NPH3 variants used (LIP motif highlighted in yellow, see text for details) and representative confocal images of leaf epidermal cells from *N. benthamiana* transiently expressing (35S promoter) RFP:NPH3 variants. Dark-adapted tobacco plants were treated with BL (∼ 11 min GFP laser). Z-stack projections of BL-treated samples are shown. Scale bars, 50 µm. White arrowheads indicate nucleus/cytoplasmic strands. All experiments were repeated at least three times with similar results. (**B**) Summary of NPH3 variants and their ability to self-associate at the plasma membrane and form cytosolic condensates. n.d. not determined. (**C**) Representative confocal images of root epidermal cells from 4-day-old light grown *Arabidopsis nph3-7* seedlings ectopically expressing constitutively condensate-forming GFP:NPH3ΔC51 and treated with 1,6-hexanediol or mock in the presence of propidium iodide (**left panel**). Note the increase in cytoplasmic GFP:NPH3ΔC51 levels upon hexanediol treatment. Scale bars: 10 μm. The **right panel** shows the number of condensates. n = 24 seedlings (untreated), n = 24 seedlings (1,6-hexanediol); untreated/1,6-hexandiol P = <0.0001 (***P < 0.001). (**D**) Ribbon plot of two NPH3 trimers crosslinked via antiparallel BTB-BTB interaction, resulting in a rotation of the second trimer by approx. 110°. Note that each trimer comprises two additional BTB domains that could be crosslinked. The corresponding side view (arrow) highlights the different orientation of the trimers.

Multivalency describes the ability to participate in more than one interaction. Deletion of the BTB domain (GFP:NPH3ΔN197) has no effect on high-affinity self-association of NPH3 at the PM (Fig. 1B,C) but prevents BL-triggered phase separation in the cytosol (Figs. 4A,B), indicating that it represents another essential NPH3 interaction valence. Collectively, BTB-independent self-association of NPH3 at the PM (dark) appears to be linked to the dynamic formation of BTB-dependent higher-order homo- or hetero-oligomers in the cytosol (BL). The BTB domain of NRL protein family members is characterized by a conserved alpha-helical extension at the C-terminus (α6-α8, highlighted in light blue in all ribbon plots and schemes of NPH3) of the core BTB fold (β-sheets B1-B3, α-helices A1-A5) (Bonchuk *et al*, 2023). BTB domains enable protein-protein interactions, including multimerization and interactions with non-BTB proteins (Stogios *et al*, 2005). Potential interaction partners of GFP:NPH3 were identified in etiolated seedlings maintained in darkness or irradiated with BL (1 µmol m^−2^s^−1^, 30 min) by IP experiments coupled with mass-spectrometry (MS) based protein identification. With the exception of 14-3-3 proteins, no polypeptides enriched in cytosolic condensates (i.e. following BL) and showing a relative abundance of ≥1% of the bait protein could be identified (Table EV1), confirming previous results (Sullivan *et al*, 2021). Considering that 14-3-3 binding is not required for condensate formation of NPH3 (Reuter *et al*, 2021), we hypothesized that NPH3 forms a single-component condensate that may contain associated proteins such as 14-3-3. This in turn implies that the BTB domain can also establish homotypic interactions. According to AF2M, NPH31′C164 (devoid of the bipartite motif) could form dimers via its BTB domain (Fig. EV4B). NPH3 could therefore be a single hub that can simultaneously homopolymerize via two distinct structural domains in its N- and C-terminus, respectively. Given the reversibility of condensate formation (Haga *et al*, 2015; Reuter *et al*, 2021), the putative BTB-BTB interaction is most likely weak and occurs spontaneously. 1,6 Hexanediol disrupts weak hydrophobic interactions and has often been used to dissolve condensates (Mosesso *et al*, 2024; Dolde *et al*, 2023). The number of GFP:NPH3ΔC51 condensates decreased significantly upon 1,6 hexanediol treatment *in vivo* (Fig. 4C), suggesting that weak hydrophobic interactions indeed play a key role in phase separation of NPH3. In this context, it is of particular importance that the BTB domains apparently dimerize antiparallel, so that the second BTB domain is rotated by approx. 110°. Consequently, crosslinking of individual NPH3 trimers via BTB-mediated antiparallel interactions could enable the formation of network-like 3D assemblies, characterized by rotated positions of the individual trimers (Figs. 4D and EV4B,C). The hypothetical antiparallel BTB-BTB interaction relies on hydrophobic interactions between helices A5 of the core BTB and α8 of the C-terminal extension (Fig. EV4B). Deletion of the C-terminal extension (NPH3ΔBTBα6-8) or helix α8 alone (NPH3ΔBTBα8) prevented NPH3 condensation (Fig. 4E). The C-terminal extension of the plant-specific BTB domain is thus crucial for polymerization of NPH3 in the cytosol. However, we cannot exclude that non-BTB proteins or BTB proteins different from NPH3 are involved in and required for the formation of cytoplasmic NPH3 assemblies.

Finally, why does disruption of the hydrophobic LIP-LIP interfaces (NPH31′LIP/NPH3_4LVQ/EA) abolish phase transition of NPH3, despite the presence of the BTB domain and self-interacting CC domain? According to AF2M, deletion of the LIP motif causes a breakdown of the symmetric right-angle conformation of the individual NPH3 monomers within the trimer/homooligomer. As a result, the BTB domains are in close contact to each other, with helices A5 and α8 of the individual monomers facing each other (Fig. EV4D). We therefore assume that the formation of antiparallel BTB-BTB interactions is almost impossible.

### Phot1-dependent dephosphorylation of NPH3 does neither require condensate formation nor PM dissociation

Based on our novel data, we addressed open questions regarding the complex changes in phosphorylation patterns of NPH3. In dark-grown seedlings, multiple residues in the N-terminal IDRs of the PM-associated NPH3 are phosphorylated independently of the photoreceptor phot1 (Kimura *et al*, 2021; Pedmale & Liscum, 2007; Tsuchida-Mayama *et al*, 2008). BL induces an immediate phosphorylation of the third-last residue of NPH3 (S744), triggering association with 14-3-3, followed by PM release of NPH3 and dephosphorylation of the N-terminal residues (Reuter *et al*, 2021; Sullivan *et al*, 2021). This light-induced and phot1-dependent dephosphorylation of NPH3 is visualized by a slight shift in electrophoretic mobility of NPH3 (Haga *et al*, 2015; Kimura *et al*, 2021; Pedmale & Liscum, 2007; Tsuchida-Mayama *et al*, 2008).

It is not known whether phot1-dependent dephosphorylation (N-terminal IDRs) requires PM dissociation and/or phase transition of NPH3 in the cytosol. Despite the inability of NPH3ΔN197, GFP:NPH3ΔBTBα8 and NPH3ΔLIP to form condensates, BL irradiation induces (i) phosphorylation of their third last residue (Fig. 5A) and (ii) PM-detachment (Fig. EV5). Additionally, (iii) a shift in electrophoretic mobility is evident, demonstrating their phot1-dependent dephosphorylation. Furthermore, all light-induced processes revert upon re-transfer to darkness (Fig. 5A), eventually leading to PM re-association of the condensate-incompetent variants (Fig. EV5). Overall, we concluded that (i) the analyzed NPH3 variants exhibit the typical properties of native NPH3, except for phase separation in the cytosol, and (ii) condensation is not required for the dephosphorylation of NPH3 observed upon phot1 activation.

**Fig. 5.**
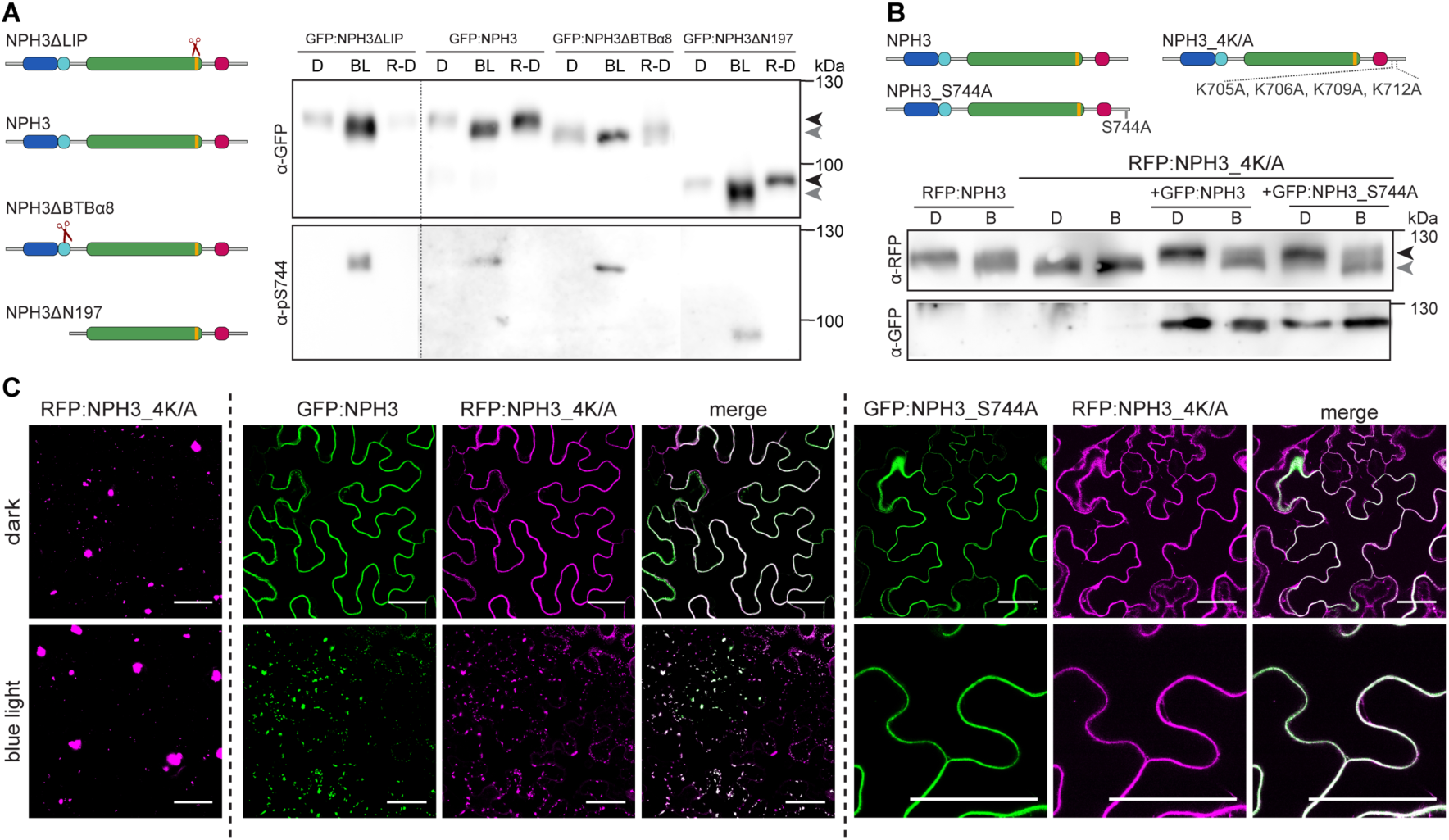
Phot1-dependent dephosphorylation of NPH3 is independent of condensate formation and PM dissociation. Black and grey arrowheads indicate the position of the phosphorylated and dephosphorylated NPH3 proteins, respectively. (**A**, **B**) Schematic representation of NPH3 variants used and immunoblot analysis of total protein extracts from *N. benthamiana* leaves transiently (co-)expressing (35S promoter) GFP/RFP:NPH3 variants. Dark-adapted tobacco plants were either maintained in darkness (D), treated with BL (10 µmol m^−2^s^−1^, 1h) (BL) or re-transferred to darkness (30 min) after irradiation (R-D). Proteins were separated on a 7.5 % SDS-PAGE gel. The dotted line indicates that NPH3ΔLIP (as compared to NPH3) was analyzed on a different gel. (**C**) Representative confocal images of leaf epidermal cells from *N. benthamiana* transiently co-expressing (35S promoter) RFP:NPH3_4K/A either with GFP:NPH3 or GFP:NPH3_S744A. Dark-adapted tobacco plants were either kept in darkness or treated with BL (∼ 11 min GFP laser). Single expression of RFP:NPH3_4K/A is shown as control. Z-stack projections of the single expression of RFP:NPH3_4K/A and of BL-treated samples are shown. Scale bars, 50 µm. All experiments were repeated at least three times with similar results.

However, the condensate-localizing NPH3_4K/A mutant is constitutively dephosphorylated in transgenic Arabidopsis seedlings (Reuter *et al*, 2021) as well as in transiently expressing *N. benthamiana* leaves (Fig. 5B). This suggests that either phot1-dependent dephosphorylation is linked to cytosolic localization of NPH3 or phot1-independent phosphorylation of NPH3-IDRs requires PM localization. We therefore relocated NPH3_4K/A to the PM in the dark by simultaneous expression with NPH3 (Fig. 5C). PM association resulted in a substantial portion of NPH3_4K/A exhibiting reduced electrophoretic mobility indicative of phosphorylation (Fig. 5B). It is worth mentioning that BL irradiation triggers dephosphorylation of NPH3_4K/A, not only when co-expressed with NPH3, but also with NPH3_S744A which keeps NPH3_4K/A attached to the PM (Fig. 5B,C). In summary, PM dissociation is dispensable for phot1-dependent dephosphorylation, whereas phot1-independent phosphorylation in the dark (N-terminal IDRs) requires PM association of NPH3. The electrophoretic mobility shift observed for the condensation-deficient NPH3 variants after re-transfer to darkness (Fig. 5A) therefore confirms their re-association with the PM.

### Condensate formation of NPH3 might be key to function

Recent studies have emphasized the critical role of light-triggered NPH3 cycling between the PM and the cytosol in phototropism (Reuter *et al,* 2021; Sullivan *et al,* 2021). However, the functional relevance of NPH3 phase separation in the cytosol remains unclear. As previously described, GFP:NPH3 was fully functional in restoring the severe impairment of hypocotyl phototropism in *nph3-7*, regardless of whether expression was driven by its native or the 35S promoter (Haga *et al*, 2015; Reuter *et al*, 2021; Sullivan *et al*, 2019; Sullivan *et al*, 2021). By contrast, ectopic expression of condensation-deficient NPH3 variants failed to complement the phenotype (Figs. 6A and EV6), suggesting that NPH3 phase separation is of importance for NPH3 function. However, GFP:NPH3ΔBTBα6-8 lacks the C-terminal extension of the BTB domain, and deletion of the LIP motif (NPH3ΔLIP) could indirectly affect BTB-dependent interactions through conformational rearrangement (Fig. EV4D). The limited function of these variants might thus alternatively be caused by loss of an essential BTB-dependent NPH3 interaction partner. A prime candidate in this regard is RPT2, which is required for hypocotyl phototropism (Haga *et al*, 2015) and has been reported to interact with NPH3 via the BTB domain. Here, the N-terminal domains of the respective proteins were tested in a yeast two hybrid assay (Inada *et al*, 2004). However, through stringent CoIP we found that the deletion of the BTB domain of NPH3 does not affect the physical association between fluorophore-tagged NPH3 and RPT2 (Fig. 6B). We therefore concluded that RPT2 associates with NPH3 *in planta* independent of its BTB domain. Considering that the typical features of NPH3, such as phosphorylation status and changes in subcellular localization, are preserved in phase transition-deficient NPH3 variants (Figs. 5A, EV5 and 6C), the most likely explanation for the impaired function is the inability to form condensates. We assume that condensation serves to sequester NPH3 in the cytosol to delay re-cycling back to the PM. Considering the reversibility of this process, the balanced temporal regulation of NPH3 cycling between the PM and the cytosol seems to be central to phototropic hypocotyl bending.

**Fig. 6.**
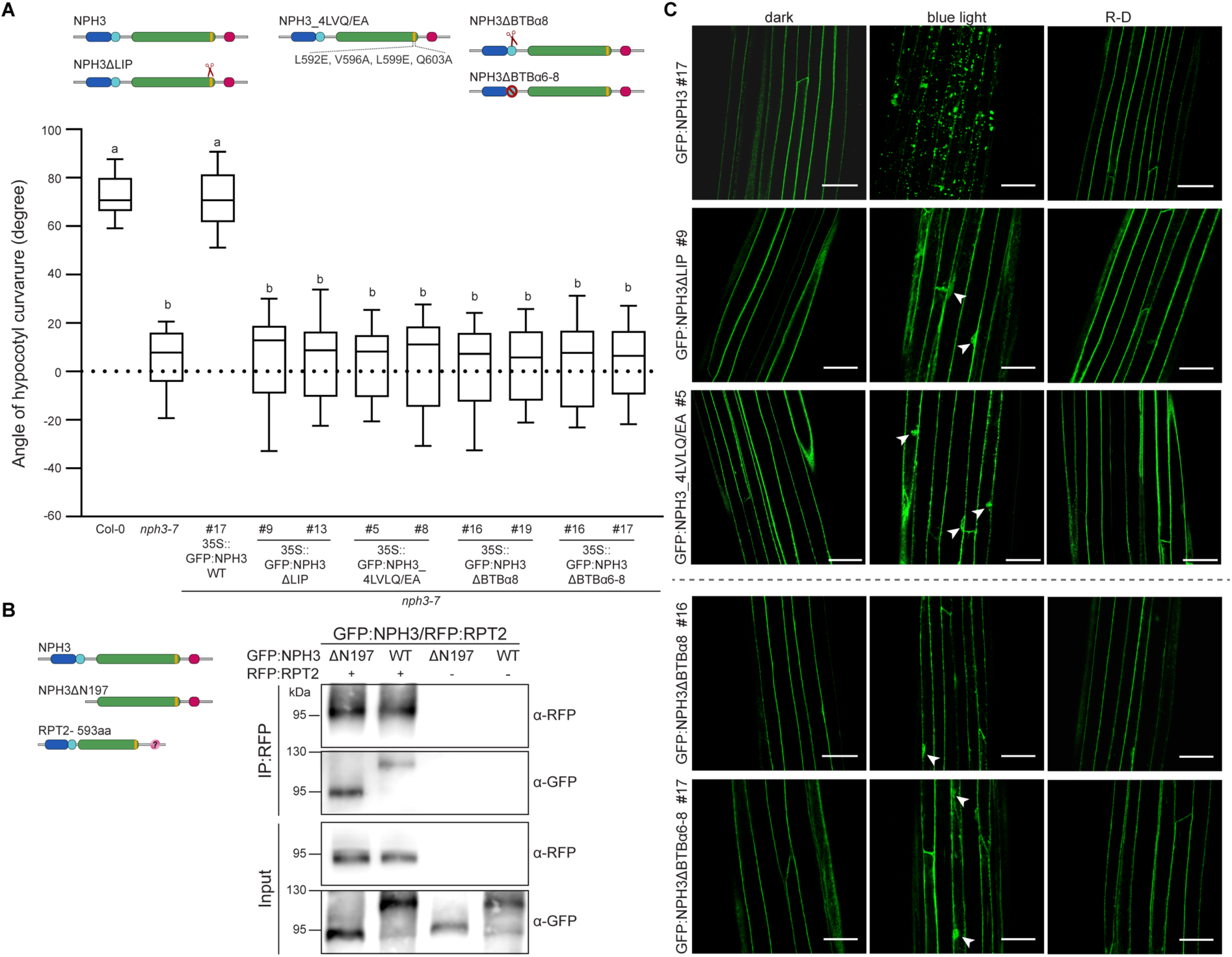
Functional relevance of NPH3 condensate formation. (**A**) Quantification of hypocotyl phototropism (mean ± SD) in etiolated *Arabidopsis nph3-7* seedlings ectopically expressing GFP:NPH3 variants and exposed to unilateral BL (0.1 µmol m^−2^ s^−1^, 24 h) (n ≥ 30 seedlings per experiment, three replicates). One way ANOVA with Tukey’s post-hoc test is shown, different letters mark statistically significant differences (p < 0.05). Centre line: median, bounds of box: minima and maxima (25^th^ and 75^th^ percentiles), whiskers: 1.5 × IQR. Exact p*-*values for all experiments are provided in the source data file. (**B**) *In vivo* interaction of GFP:NPH3 variants transiently co-expressed (35S promoter) with RFP:RPT2 in *N. benthamiana* leaves (dark-adapted). The crude protein extract was immunoprecipitated using RFP beads. Immunodetection of input and immunoprecipitated (IP:RFP) proteins is shown. Experiments were performed at least two times with similar results. (**C**) Representative confocal microscopy images of hypocotyl cells from 3-day-old etiolated *Arabidopsis* seedlings shown in (A), and either maintained in darkness (dark), treated with BL (1 µmol m^−2^s^−1^) for 1 hour or re-transferred to darkness (1 h) after irradiation (R-D). Note that when analyzing the three transgenic lines shown below the dashed line, GFP laser power was increased due to a weaker expression of the transgenes (Fig. EV6). Z-stack projections of BL-treated samples are shown. Scale bars, 50 µm. Experiments were performed at least two times with similar results. Arrowheads indicate nucleus.

## Discussion

Our data provide novel insights into molecular signatures of NPH3 defining its function in BL-induced phototropic hypocotyl bending. We applied a combination of live-cell imaging, deep learning-based structure prediction, biochemical, genetic and physiological approaches to uncover unique, overlapping and cooperative roles of NPH3 motifs essential for self-association at the PM and/or the formation of multivalency-driven condensates in the cytosol. Members of the plant NRL protein family are mostly characterized by three domains, the N-terminal BTB-, the central NPH3 domain and the C-terminal CC domain, all of which are known since decades (Motchoulski & Liscum, 1999). However, our understanding of the structure-function relationships of NRL’s is quite limited.

*In vivo* protein-protein interaction assays, combined with confocal imaging of *N. benthamiana* leaf epidermis cells expressing NPH3 variants revealed a crucial role of the CC domain in mediating high-affinity self-association of NPH3 at the PM (Fig. 2A,B). Although deletion/destabilization of the CC (NPH3ΔC104/NPH3_WLM/E) substantially impaired the interaction strength, weak homo-oligomerization and, above all, condensate formation could still be observed (Figs. 2A,B,E and 4A). AF2M/AF3 predicted the formation of NPH3 homooligomers (≥3), most likely trimers and provided previously unexpected molecular insights that eventually led to the discovery of a bipartite C-terminal self-association motif. In fact, the trimeric NPH3 complex appears to be stabilized by (i) the CC superhelices and (ii) a helical bundle formed by a yet unrecognized upstream structural feature, the LIP motif (Fig. 2C). Experimental validation confirmed homo-oligomerization of the CC domain (Fig. EV3C) and emphasized the role of the LIP motif in establishing low-affinity homomeric association (Figs. 2B,E and EV2B). The LIP motif localizes to the C-terminal end of the NPH3 domain and has received little attention to date. However, considering that about one third of the 33 members of the NRL protein family are devoid of a CC domain (Pedmale *et al*, 2010), while the LIP motif is highly conserved (Fig. EV3B), LIP-mediated self-association with low affinity is most likely a functionally significant feature. This is supported by the – exclusively LIP-mediated - trimer formation of NPH3ΔC104 predicted by AF2M (Fig. EV3D).

Since NPH3 homo-oligomerization is mediated by a bipartite structured motif, we examined whether its individual parts serve distinct functions. Self-association linked to relocation appears to be a unique feature of the CC domain and could potentially alter the subcellular localization of cytosolic NRLs through heteromeric complex formation with a PM-associated member. The second molecular function of the bipartite motif relates to PM association of NPH3. Here, we demonstrated that homo-oligomerization of NPH3 in addition to its electrostatic interaction with phospholipids (Reuter *et al*, 2021) is essential for PM recruitment (Fig. 3B-D). A comparable scenario has been described for nanodomain-organizing REMORINs (Martinez *et al*, 2019). Homo-oligomerization of NPH3 could enable folding/exposure of the amphipathic helix, which is essential for phospholipid binding, or alternatively increase avidity towards phospholipids. In the context of PM attachment, the two parts obviously have an unequally redundant function. While the CC domain mediates strong PM association, the LIP motif facilitates weak PM binding when the CC domain is destabilized (NPH3_WLM/E) (Fig. 3 D,E). Thirdly, the C-terminal bipartite self-association motif contributes to the higher-order oligomeric states of phase-separated NPH3 (NPH3ΔC164) (Fig. 4A). Notably, only the LIP motif (low affinity) is of importance here and acts cooperatively with the N-terminal BTB domain (weak interaction) in condensate formation (Fig. 4A), consistent with the finding that membrane-less assemblies can be formed by an ensemble of low-affinity molecular interactions (Boija *et al*, 2018). Since interaction partners of NPH3 that are essential for condensate formation have not yet been identified (Table EV1) (Sullivan *et al*, 2021), we suspected NPH3 to form a single-component condensate. Consequently, LIP-mediated self-association of NPH3 at the PM appears to be coupled to weak BTB-mediated homo-oligomerization in the cytosol, facilitating the formation of higher-order complexes. Based on the AF2M prediction, the BTB domain (NPH3ΔC164 without the bipartite C-terminal motif) may facilitate antiparallel dimerization (Fig. EV4B). This process would lead to the formation of crosslinked and rotated trimers, ultimately resulting in a 3D network of polymerized trimers in the cytosol (Figs. 4D and EV4C). However, the 3D arrangement of the individual trimers also influences the positioning of the phospholipid-binding motif, potentially hindering effective binding on the 2D space of the PM. We therefore propose that this affine phospholipid binding restricts higher-order 3D oligomerization via weak BTB-BTB interactions at the PM. In summary, NPH3 acts as a bistable switch regulated by its subcellular localization. Here, a strict C-terminal trimerization at the PM is transformed into a co-operative bivalent (LIP and BTB) homo-oligomerization in the 3D space of the cytosol.

During eukaryotic evolution, BTB domains progressively adapted to fulfill diverse cellular functions (Emenecker *et al*, 2021; Field *et al*, 2023). NRL family members are plant-specific BTB domain proteins and feature a conserved extension at the C-terminus of the BTB domain (α6 - α8) (Bonchuk *et al*, 2023), which is crucial for condensate assembly of NPH3 (Fig. 4E). In addition, the NRL BTB domains adopted an N-terminal extension (Bonchuk *et al*, 2023) with high structural variability among NRL family members. These variations could affect the ability, strength and orientation of BTB homo-oligomerization. It is worth noting that NPH3 may form part of a CULLIN3 RING E3 UBIQUITIN LIGASE (CRL3) (Roberts *et al*, 2011). Several – non NRL - plant BTB domain proteins act as substrate adapters in CRL3s with the BTB domain responsible for binding to the scaffold CULLIN3 (CUL3) (Ban & Estelle, 2021). BTB dimerization by apposition of two monomers in the same orientation is important for the function of many CRL3 orthologues in humans/yeast. Such BTB dimer formation is unlikely for NPH3 (Bonchuk *et al*, 2023) (Fig. EV4B) and CUL3 was not identified as an NPH3 interaction partner in transgenic plants (EV Table1) (Sullivan *et al*, 2021). Clearly, further biochemical work is needed to elucidate the possible involvement of NRL BTB domains in protein ubiquitylation. Variations in the N-terminal extension of NRL BTB domains could furthermore impact on homo-oligomerization of NRLs in general (CC/LIP *versus* BTB). NRL family members lacking a stringent CC domain might therefore rely either on the BTB domain or the LIP motif for homo-oligomerization, depending on their respective affinities. In this context, NPY8/NRL5 which localizes close to the PM and belongs to a different NRL clade, was recently described to homomerize via its BTB domain based on *in vitro* pull-down assays (Upadhyay-Tiwari *et al*, 2024). Conversely, both the homo-oligomerization of NPH3 at the PM (Fig. 1C) and its interaction with RPT2 (Fig. 6B) occurs independently of its BTB domain *in planta*. Interestingly, AF2M predicts NPY8/NRL5 to form either a BTB-mediated dimer or alternatively a LIP-mediated trimer. It is therefore of importance to validate these scenarios experimentally, taking into account possible changes in subcellular localization.

The use of biomolecular condensates as a means of spatial and temporal regulation is a ubiquitous phenomenon observed in all kingdoms of life. Plant biomolecular condensates are increasingly recognized as playing important roles in plant development and environmental responses. Prominent examples encompass flowering, auxin signaling, seed germination, RNA processing and immunity (Emenecker *et al*, 2021; Field *et al*, 2023). Reversible protein assemblies are usually drivers of either recruitment or sequestration. Sequestration of the auxin response factor ARF19 into cytosolic condensates regulates auxin transcriptional responses in a cell-specific manner by keeping the transcription factor away from the nucleus (Powers *et al*, 2019). Regarding NPH3, transitioning into a condensed state likely sequesters the protein in the cytosol, thereby regulating its rate of re-association with the PM, which can be observed not only upon re-transfer to darkness (Reuter *et al*, 2021) but also upon prolonged irradiation (Haga *et al*, 2015). Mutations that disrupt NPH3 condensates almost completely abolished phototropic hypocotyl bending (Fig. 6A), suggesting that a precise dwell time of NPH3 in the cytosol is critical for function. However, we cannot rule out the possibility that the limited function of these variants is due to loss of an essential interaction that requires the C-terminal extension of the BTB domain of NPH3. Such an *in vivo* interaction partner has, however, not yet been identified (EV Table1) (Sullivan *et al*, 2021). Overall, our data suggest that phase separation provides a mechanism to regulate cycling of NPH3 between the PM and the cytosol, particularly temporally. The elucidation of a putative important function of NPH3 at the PM, which must be modified in a transient and temporally tightly controlled manner to trigger phototropic hypocotyl bending, represents a major challenge for future research.

## Material and Methods

### Plant materials, transformation and growth conditions

Seeds of *Arabidopsis thaliana* [ecotype Columbia-0 (Col-0), *nph3-7* (SALK_110039, Col-0 background)] were obtained from the Nottingham Arabidopsis Stock Centre, and the *nph3-7* transgenic lines expressing GFP:NPH3 or GFP:NPH31′C51 have been described recently (Reuter *et al*, 2021). Stable transformation of *nph3-7* followed standard procedures.

Unless otherwise stated, seeds were surface sterilized and plated on solid half-strength Murashige and Skoog (MS) medium (pH 5.8) containing 1 % sucrose (w/v). Following stratification in the dark for 48-72 h at 4°C, seeds were exposed to fluorescent white light for 4 h. Seedlings were then grown in darkness for 68 h at 20°C. Subsequently, the etiolated seedlings were either kept in darkness or irradiated with BL [overhead BL (1 μmol m^−2^s^−1^) for up to 1 hour or, alternatively, GFP-laser treatment (488 nm) for up to 11 min during confocal observation, as specified in the Figure legends]. Independent experiments were carried out at least in triplicates. Representative images are presented.

*Agrobacterium*-mediated transient transformation of 3-4 weeks old *Nicotiana benthamiana* plants was performed as described previously (Blatt & Grefen, 2014). *Agrobacterium tumefaciens* strain GV3101, transformed with a binary vector of interest, was resuspended in infiltration solution (10 mM MES, pH 5.6, 10 mM MgCl_2,_ 150 µM acetosyringone) at an OD_600_ of 0.1 – 0.2 and syringe-infiltrated into the abaxial epidermis of *N. benthamiana* leaves. For co-transformation, a 1:1 mixture was used, sometimes combined with co-introduction of a suppressor of post-transcriptional gene silencing [35S::P19 (Win & Kamoun, 2004), OD_600_: 0.01 – 0.05]. Freshly transformed tobacco plants were kept under constant light for 24 h, subsequently transferred to darkness for 17 h (dark adaptation) and finally either kept in darkness (D) or BL-irradiated [overhead BL (10 μmol m^−2^s^−1^) for up to 1 hour or, alternatively, the GFP-laser treatment (488 nm) for up to 11 min during confocal inspection of abaxial leaf epidermis cells, as specified in the Figure legends]. Independent experiments were carried out at least in triplicates. Representative images are presented.

### Cloning procedures

Cloning of N-terminally fluorophore-tagged NPH3 variants (GFP and/or RFP) into the destination vectors pB7WGR2 and/or pH7WGF2 (Karimi *et al*, 2007) for stable transformation or transient overexpression followed standard GATEWAY™ procedures. The 35S-driven NPH3 variants GFP/RFP:NPH3, GFP/RFP:NPH3_S744A, GFP/RFP:NPH3ΔC51 and GFP/RFP:NPH3_4K/A transformation vectors have been described before (Reuter *et al*, 2021). Transgenic plants were selected based on the hygromycin resistance conferred by pH7WGF2 and homozygous lines were established.

Site-directed mutagenesis was performed by PCR, while deletions were generated by primer extension PCR. PCR products and products of mutagenesis were verified by sequencing. A complete list of oligonucleotides used in this study for PCR is provided in EV Table 2.

### Expression and purification of proteins

For bacterial expression of the Arabidopsis C-terminal NPH3 variants N-terminally tagged with (His)_6_-Maltose Binding Protein (MBP), the corresponding cDNA was amplified by PCR and cloned into the expression vector pDEST-HisMBP (Addgene plasmid #11085) using standard GATEWAY^TM^ procedures. Recombinant NPH3 proteins were expressed using *Escherichia coli* BL21(DE3) host strain. Transformed *E.coli* BL21(DE3) was grown in liquid 2 x yeast extract and tryptone (2YT) medium containing ampicillin (100 μg/ml) at 37°C until an OD_600_ of 0.6. Protein expression was induced by adding IPTG to a final concentration of 0.5 mM. Following overnight growth at 16°C, bacteria were harvested by centrifugation (4000 rpm, 30 min, 4 °C). The pellet was frozen in liquid nitrogen. Following thawing on ice the cells were resuspended in bacterial protein extraction reagent B-PER (5 ml/g fresh weight) (Thermo Scientific, Waltham, USA) supplemented with Protease Inhibitor Mix-B (Serva, Heidelberg, Germany). Purification under native conditions was done by using the cleared BL21(DE3) lysate and Ni^2+^-NTA agarose (Qiagen, Hilden, Germany) according to the manufacturer’s protocol.

### Size Exclusion Chromatography (SEC)

SEC was performed using the Superdex 200 Increase 10/300 GL column in combination with the ÄKTA^TM^ pure chromatography system (Cytiva, Marlborough, USA). 200 μl of purified recombinant protein (approx. 2 mg/ml) was loaded onto the PBS-equilibrated (10 mM phosphate buffer, 140 mM NaCl, pH 7.4) column using a flow rate of 0.1 ml/min (UNICORN^TM^ 7.0 program, Cytiva), and collected in 0.5 ml fractions using a 96-well fraction collector (Biozym, Oldendorf, Germany). The collected fractions were subjected to immunoblot analysis. The molecular weight of the recombinant NPH3 proteins was calibrated using standard size protein markers [High Molecular Weight Gel Filtration Calibration Kit (Cytiva): thyroglobulin (669 kDa, 5 mg/ml), Ferritin (440 kDa, 0.3 mg/ml), Aldolase (158 kDa, 4 mg/ml), Conalbumin (75 kDa, 3 mg/ml), Albumin (44 kDa, 4 mg/ml) and Chymotrypsinogen A (25 kDa, 3 mg/ml)].

### Preparation of microsomal membranes

Microsomal membrane fractions were prepared from transiently transformed and dark-adapted *N. benthamiana* leaves. Tissue was homogenized with 3 ml of ice-cold homogenization buffer per g fresh weight [50 mM Hepes, pH 7.8, 500 mM sucrose, 1 % PVP-40, 3 mM dithiothreitol (DTT), 3 mM EDTA, supplemented with Complete Protease Inhibitor Mixture (Roche, Basel, Switzerland) and Phosphatase Inhibitor Mix 1 (Serva)]. The homogenate was centrifuged at 10,000 × g for 20 min at 4°C. The supernatant was filtered through MiraCloth and subsequently centrifuged at 100,000 × g for 45 min at 4°C. The microsomal pellet was resuspended in ice-cold 5 mM Tris/MES, pH 6.5, 330 mM sucrose, 2 mM DTT, supplemented with Complete Protease Inhibitor Mixture (Roche) and Phosphatase Inhibitor Mix 1 (Serva). All the above steps were performed under red safe light conditions.

### Co-Immunoprecipitation and MS analysis

*In vivo* interaction of fluorophore-tagged NPH3 variants was tested by using total plant protein extracts obtained from transiently transformed *N. benthamiana* leaves according to (Albert *et al*, 2015) with slight modifications. Dark-adapted tobacco plants were either kept in darkness or treated with overhead BL (10 μmol m^−2^s^−1^) for 1 h. Approximately 500 mg of leaf tissue was ground thoroughly under red safe light illumination and suspended in solubilization buffer [50 mM Tris, pH 7.5, 150 mM NaCl, 1 mM EDTA, 0.75% Triton X-100 supplemented with Complete Protease Inhibitor Mixture (Roche) and Phosphatase Inhibitor Mix 1 (Serva)]. After 1 h incubation on an overhead rotor at 4 °C, cell debris-removed supernatant was incubated with equilibrated 20 μl GFP-Trap® or RFP-Trap® agarose beads (ChromoTek and Proteintech Company, Planegg-Martinsried, Germany) for 1 h in the cold room with overhead rotation. The beads were washed three times with solubilization buffer containing 0.1% Triton X-100, followed by a final washing step using 50 mM Tris, pH 7.5, 150 mM NaCl. Finally, proteins were eluted by adding 40 µl of 2 × SDS sample buffer (100 mM Tris-HCl, pH 6.8, 20% glycerol, 2% β-Mercaptoethanol, 4% SDS, 0.02% bromophenol blue) and boiling at 95 °C for 10 min. The denatured proteins were subsequently separated by SDS-PAGE. Arabidopsis *nph3-7* ectopically expressing GFP:NPH3 and, as control, GFP (UBQ10 promoter) were sown on half-strength MS plates and grown in the dark for 3 days. Subsequently, the etiolated seedlings were either kept in darkness or treated with overhead BL (1 μmolm^−2^ s^−1^) for 30min. Three grams of plant tissue were used under red safe light illumination for immunoprecipitation essentially as described above. Cell debris-removed supernatants were incubated with 50 μl GFP-Trap beads (ChromoTek) for 3 h in the cold room with mild rotation. The washing steps were supplemented by a final wash (3x) with 20mM ammoniumbicarbonate buffer. On-beads digestion with trypsin, peptide purification with Phoenix kit (PreOmics) and liquid chromatography-MS/MS analysis (Proxeon Easy-nLC coupled to an QExactiveHF-X mass spectrometer, Thermo) were performed by the Proteome Center Tübingen, University of Tübingen, essentially as described (Schmitt *et al*, 2021), except that peptides with a resolution of 45,000 were selected for MS/MS sequencing.

### Protein quantification, SDS-PAGE and Western Blotting

Protein concentration was determined using ROTI® Nanoquant (Carl Roth, Karlsruhe, Germany) according to the manufacturer’s protocol.

SDS-PAGE, Western blotting and immunodetection followed standard procedures. Total proteins were extracted from transiently transformed *N. benthamiana* leaves (2 leaf discs) by directly grinding in 100 μl 2 × SDS sample buffer under red safe light illumination.

Total protein extracts from 3-day-old etiolated Arabidopsis seedlings (150 seedlings) were prepared by grinding the frozen tissue in ice-cold extraction buffer (20 mM Tris, pH 7.0, 150 mM NaCl, 1 mM EDTA, 1 % Triton X-100, 0.1 % SDS, 5 mM DTT) supplemented with Complete Protease Inhibitor Mixture (Roche) and Phosphatase Inhibitor Mix I (Serva). After 10 minutes incubation on ice, cell debris-removed supernatants were used for immunodetection. Chemiluminescence was detected using an Amersham ImageQuant 800 (Cytiva) system. Images were processed with Adobe Photoshop CS5 (Adobe Inc., San José, CA, USA) for adjustment of brightness and contrast. Chemiluminescence detection was performed using the ECL™ Prime Western Blot Detection Reagent (Cytiva).

The following antibodies were used in this study: anti-NPH3-S744P (Reuter *et al*, 2021) (1:500), anti-NPH3 (Reuter *et al*, 2021) (1:1000), anti-GFP (1:1000, Thermo Scientific, catalog number: A-11122, lot: 2477546), anti-RFP 5F8 (1:1000, ChromoTek and Proteintech Company, catalog number: 5f8-150, lot: 104272-05), anti-His_6_ (1:2000, Roche/Merck, Darmstadt, Germany, catalog no: 11965085001).

### Hypocotyl Phototropism analysis

Phototropism was performed using *A. thaliana* seedlings which were grown in the dark for 72 h on vertically oriented solid half-strength MS plates containing 1 % sucrose. Etiolated seedlings were then transferred to a LED chamber and illuminated with unilateral BL (0.1 μmol m^−2^ s^−1^) for 24 h. Plates were scanned, and the inner hypocotyl angle was measured for each seedling using ImageJ (Schindelin *et al*, 2012). The curvature angle was calculated as the difference between 180° and the measured value. For each transgenic line three biological replicates (*n*≥30 seedlings per experiment) were performed, alongside with the appropriate controls (Col-0, *nph3-7*).

### AlphaFold Predictions

To predict protein and protein complex structures we used the AlphaFold2-Multimer program developed by Google DeepMind (Evans *et al*, 2022; Jumper *et al*, 2021) through a subscription to the Google Colab v1.5.5 (Mirdita *et al*, 2022) (https://colab.research.google.com/github/sokrypton/ColabFold/blob/main/AlphaFold2.ipynb) following guidelines on the document. As an alternative, AF3 predictions (developed by Google DeepMind and Isomorphic Labs) (Abramson *et al*, 2024) were generated via the newly launched AlphaFold server (https://alphafoldserver.com). Models were colored in PyMol (The PyMOL Molecular Graphics System, Version 2.5.8 Schrödinger, LLC). Where indicated, predictions are colored according to the AF2 produced per-residue confidence metric called the local distance difference test (pLDDT) (Elfmann & Stülke, 2023; Evans *et al*, 2022) via visualization in PyMol.

### Confocal microscopy

Live-cell imaging was performed using the Leica TCS SP8 (upright) confocal laser scanning microscope. Imaging was done by using a 63x/1.20 water immersion objective. For excitation and emission of fluorophores, the following laser settings were used: GFP, excitation 488 nm, emission 505-530 nm; RFP, excitation 558 nm, emission 600-630 nm. All CLSM images in a single experiment were captured with the same settings using the Leica Confocal Software.

For simultaneous imaging of samples expressing two fluorophores, sequential scanning mode was used supported by Leica Confocal Software. All experiments were repeated at least three times. Both the focal plane images and Z-stack maximum projection images were processed using Leica Application Suite X 3.7.4.23463.

For *in planta* FRET-FLIM analyses, *N. benthamiana* plants were analyzed 2 days after transient transformation using an SP8 confocal laser scanning microscope (Leica Microsystems), which was equipped with the Leica Microsystems Application Suite software and a FastFLIM upgrade from PicoQuant (Sepia multichannel picosecond diode laser, PicoQuant TimeHarp 260 TCSPC module, and Picosecond Event Timer). The measurements were performed according to (Ladwig *et al*, 2015) and (Glöckner *et al*, 2022) with some adjustments. Imaging was performed using a 63x/1.20 water-immersion objective, focusing on the PM of abaxial epidermal cells. Prior to the FRET-FLIM measurements, the presence of the fluorophore-fusions was confirmed by using 488 or 561 nm lasers for excitation of GFP or RFP, respectively. The donor lifetime was determined in donor-only expressing cells or cells with the indicated combinations using a pulsed laser as an excitation light source at 470 nm and a repetition rate of 40 MHz. The acquisition was performed until 500 photons were reached in the brightest pixel, with a resolution of 256 x 256 pixels and a pixel dwell time of ∼ 20 µs. The corresponding emission was detected with a HyD SMD from 500 nm to 550 nm by TCSPC.

For condensate analyses in vivo, 4-day-old seedlings grown on half-strength MS plates supplemented with 1 % sucrose were transferred on a microscopy slide with either water or 10% 1,6-hexanediol (Merck, 240117), supplemented with 3 µg/ml propidium iodide for viability test, immediately before imaging. Root epidermal cells were analyzed 3 min after treatment. For counting the number of puncta, a threshold was set for each image, the images were converted to binary images, and masked. Region of Interests with uniform fluorescence were selected and the number of particles was counted using the “Analyze Particles” function in Fiji with 0.05/0.1 to 8/10 μm^2^ as size range and 0 to 1 circularity. The number of puncta was then normalized for the area of 10000 μm2.

### Data analysis

For all semi-quantitative CoIPs, the protein band intensities of GFP and RFP from IP:GFP were determined by Western blot quantification using ImageJ software as described by Hossein Davarinejad (http://www.yorku.ca/yisheng/Internal/Protocols/ImageJ.pdf). As a next step, the ratio of RFP band intensity to the corresponding GFP band intensity was determined. The values given in Fig. 2B were normalized to NPH3WT/NPH3ΔC104 (dark; left panel) and NPH3_WLM_E/NPH3ΔC51 (dark; right panel). Likewise, the values in Fig. EV2B were normalized to NPH3ΔC51/NPH3ΔC104 (dark).

FRET-FLIM: The SymPhoTime 64 software (Vers. 2.9) was employed for data processing. Iterative reconvolution was used to calculate the FLTs, i.e. the instrument response function was convolved with a bi-exponential decay function, for both the donor-only expressing cells and the indicated combinations. Only the range between 1.3 ns and 20 ns was used for fitting. To calculate the percentage of IPS in FRET-FLIM measurements we made use of custom-made R-Scripts and set adjusted thresholds according to (Bücherl *et al*, 2013). The scripts will be made accessible upon completion of the peer-review process or upon request. The photon counts per pixel must be at least 250. For each pixel that met the aforementioned criterion, the FLT was derived by intensity-weighted averaging and the values were exported from the SymPhoTime 64 software. For further processing, pixels with FLTs below 1.5 and above 2.7 were excluded to avoid false positive or negative interactions. Subsequently, the mean FLT of all donor-only samples were determined. Based on this value and the threshold of 10 % FRET efficiency (Glöckner *et al*, 2022), the limit of IPS was defined. According to this definition, all cells were analyzed and only pixels with FLTs below the interaction threshold were collected as IPS. The ratio between the IPS and the total amount of selected pixels represents the value of IPS in the respective graph.

IP-MS: Processing of the data was conducted using MaxQuant software (vs 2.2.0.0.). The spectra were searched against an *A. thaliana* UniProt database. Raw data processing was done with 1% false discovery rate setting. Two individual biological replicates were performed and mitochondrial, chloroplastic and peroxisomal proteins were omitted from the list of GFP-NPH3 interaction partners. Only proteins identified in two biological replicates from at least two peptides were retained for analysis. Proteins identified in IPs from GFP-expressing seedlings (≥ 3 peptides or number of peptides close to the bait) were considered contaminants. Protein Intensity-Based Absolute Quantification values (iBAQ) were converted to normalized iBAQ of the bait protein (GFP:NPH3). Proteins showing at least a twofold change in iBAQ values following blue light irradiation were identified (Table EV1).

The number of condensates from seedlings expressing GFP:NPH3ΔC51 was analyzed using ImageJ software (see confocal microscopy). Graphical and statistical analysis (two-tailed unpaired *t-*test with unequal variance) were performed using the software GraphPad Prism. Error bars represent SD.

The angles of phototropic curvature of all analyzed Arabidopsis genotypes (≥ 30 seedlings per genotype per replicate) were measured using ImageJ software. Graphical analysis was performed using the software GraphPad Prism. Statistical significance of the data was assessed using one-way ANOVA followed by *post hoc* Tukey multiple comparison test (*P* < 0.05). Error bars represent SD.

## Acknowledgements

We extend our gratitude to Nathalie Zgoda for her contributions to data analysis, and to Franziska Lanz and Ricarda Furkert for their assistance with the analysis of the recombinant C-terminal NPH3 fusion proteins. We also appreciate the SEC support provided by Chenlei Hua and Lisha Zhang, and the SP8 support from Sandra Richter. Lastly, we thank John Christie and Thorsten Nürnberger for their valuable comments on the manuscript.

## Funding

This work was supported by the German Research Foundation (DFG) with a grant to CO (CRC 1101-B09) and grants for scientific equipment (INST 37/819-1 FUGG, INST 37/991-1 FUGG).

## Author contributions

CO conceived the project. PM, LRe and CO designed experiments. PM, LRe, LRo, AF, YIY, TS, ID-B, JK and AB performed experiments. CO performed AF2/AF2M/AF3 predictions. PM, LRe, LRo, TS, ID-B and CO analyzed data. The manuscript was principally written by CO with comments from all authors.

## Competing interests

The authors declare that they have no competing interests.

## Data and materials availability

All data needed to evaluate the conclusions in the paper are present in the paper and/or the Supplementary Materials. Requests for material should be submitted to Claudia Oecking.

---

**Figure EV1.**
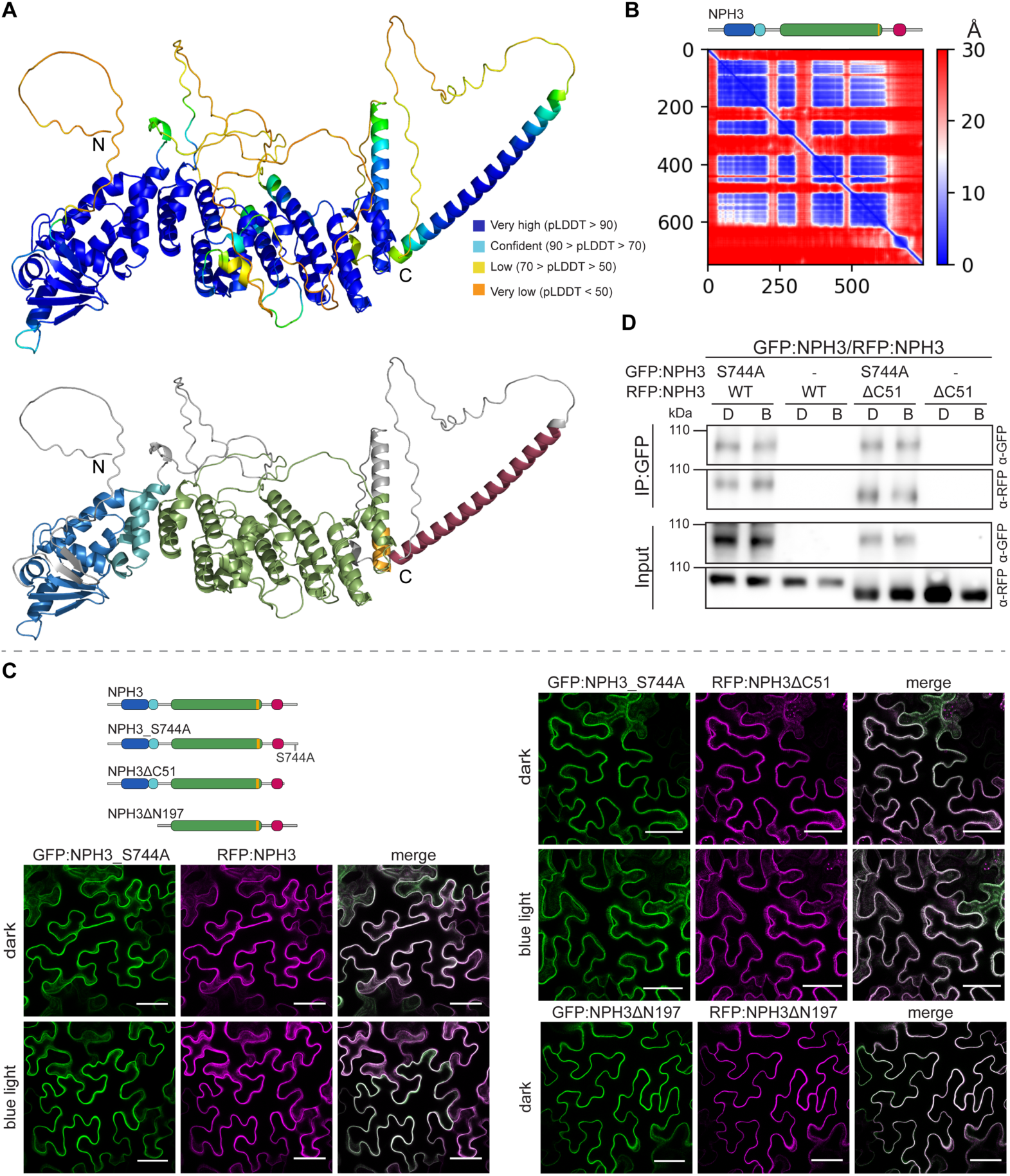
Self-association of NPH3 is independent of its subcellular localization. (**A**) Protein structure of NPH3 predicted by AF2. Color-coding of the ribbon plot is based either on the ‘per residue confidence score’ (pLDDT, **upper panel**) or on the schematic domain representation (Fig. 1A) (**lower panel**). N, N-terminus; C, C-terminus. (**B**) PAE plot of the NPH3 structure shown in (A). (**C**) Representative confocal images of leaf epidermal cells from *N. benthamiana* transiently co-expressing (35S promoter) GFP/RFP:NPH3 variants. Dark-adapted tobacco plants were either kept in darkness or treated with BL (∼ 11 min GFP laser). Z-stack projections are shown, except for GFP:NPH3ΔN197/RFP:NPH3ΔN197. Scale bars, 50 µm. These experiments were repeated at least three times with similar results. (**D**) *In vivo* interaction of GFP/RFP:NPH3 variants transiently co-expressed (35S promoter) in *N. benthamiana* leaves. Dark-adapted tobacco plants were either kept in darkness (D) or treated with BL (B) (10 µmol m^−2^s^−1^, 1h). The crude protein extract was immunoprecipitated using GFP beads. Immunodetection of input and immunoprecipitated proteins (IP:GFP) is shown. Experiments were performed at least two times with similar results.

**Figure EV2.**
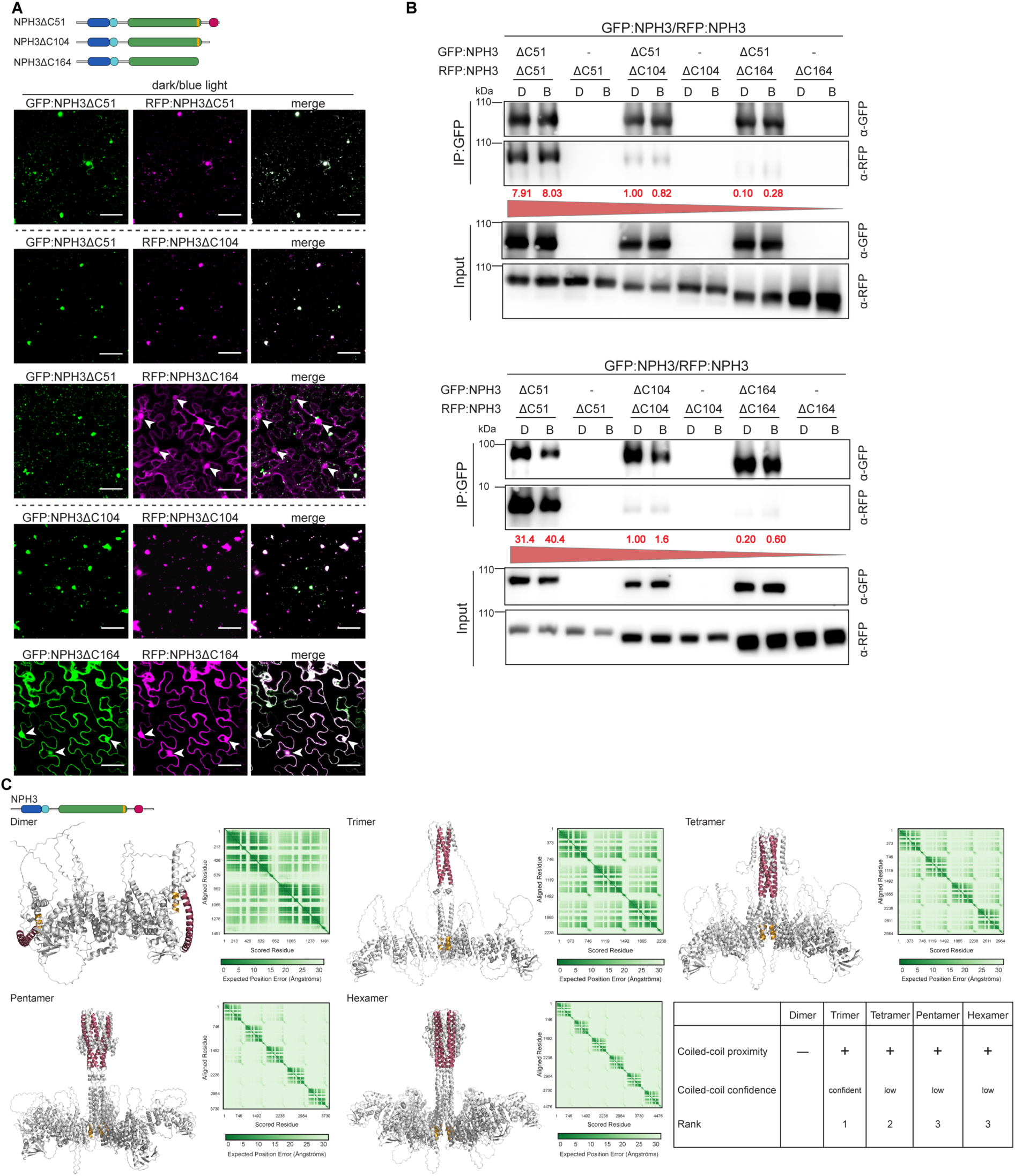
The bipartite C-terminal self-association motif of NPH3 consists of the CC domain and the LIP motif. (**A**) Schematic representation of NPH3 variants used and representative confocal images of leaf epidermal cells from *N. benthamiana* transiently co-expressing (35S promoter) GFP/RFP:NPH3 variants. Dark-adapted tobacco plants were either kept in darkness or treated with BL (∼ 11 min GFP laser). Z-stack projections are shown. Scale bars, 50 µm. Experiments were performed at least three times with similar results. White arrowheads indicate nucleus. (**B**) *In vivo* interaction of GFP/RFP:NPH3 variants transiently co-expressed (35S promoter) in *N. benthamiana* leaves. Dark-adapted tobacco plants were either kept in darkness (D) or treated with BL (B) (10 µmol m^−2^s^−1^, 1h). The crude protein extract was immunoprecipitated using GFP beads. Immunodetection of input and immunoprecipitated proteins (IP:GFP) is shown. Experiments were performed at least three times with similar results. Values in red: normalized RFP/GFP intensity ratio. (**C**) Ribbon plot and PAE plot of NPH3 homooligomers predicted by AF3. The CC domain and the LIP motif are color-coded according to the schematic domain representation (Fig. 1A). A comparative analysis of the characteristics of the respective homooligomers is given.

**Figure EV3.**
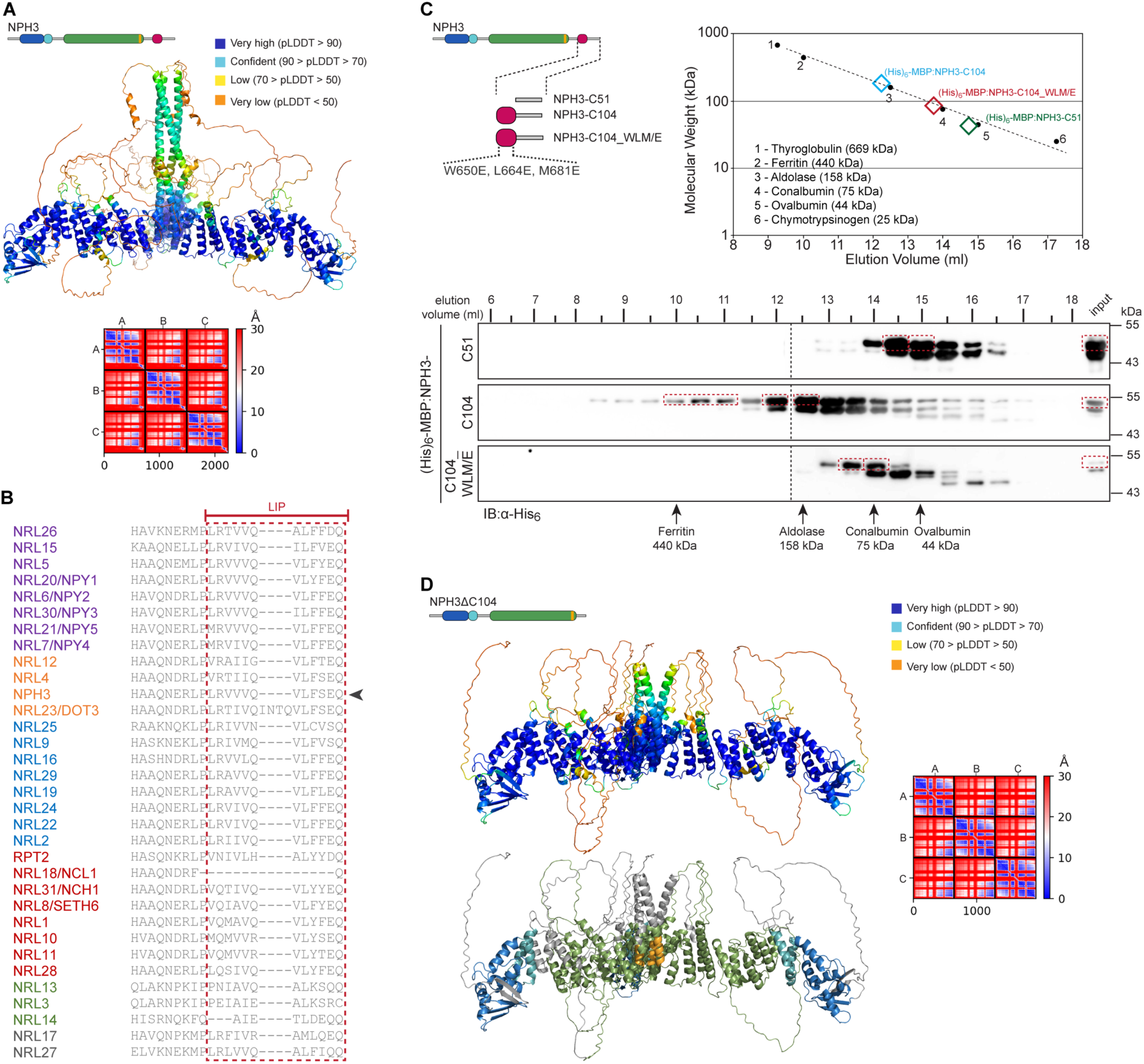
Homo-oligomerization of the CC domain and LIP motif conservation. (**A**) Ribbon plot of NPH3 trimer predicted by AF2M and color-coded according to the pLDDT score (**upper panel**). The corresponding PAE plot is shown in the **lower panel**. (**B**) Amino acid alignment of the *Arabidopsis* NRL protein family focusing on the residues constituting the LIP motif in NPH3 (red dotted box). (**C**) Schematic representation of C-terminal NPH3 fragments **(left panel)** and size exclusion chromatography **(right panel)** of the purified fragments tagged with (His)_6_-Maltose Binding Protein (MBP). Elution of molecular mass marker proteins (1-6) is indicated within the graph. Molecular weights of the proteins are given in logarithmic scale (Y-axis). The α-His immunological analysis of fractions obtained via size exclusion chromatography is shown in the **lower panel**. The red dotted boxes highlight the purified non-degraded fusion protein in the input as well as in the elution fractions. Arrows mark the elution peaks of the marker proteins with the indicated molecular mass. (**D**) Ribbon plot of the NPH3ΔC104 trimer predicted by AF2M and color-coded according to the pLDDT score (**upper left panel**) or the schematic domain representation (Fig. 1A) (**lower left panel**). The corresponding PAE plot is shown in the **right panel**.

**Figure EV4.**
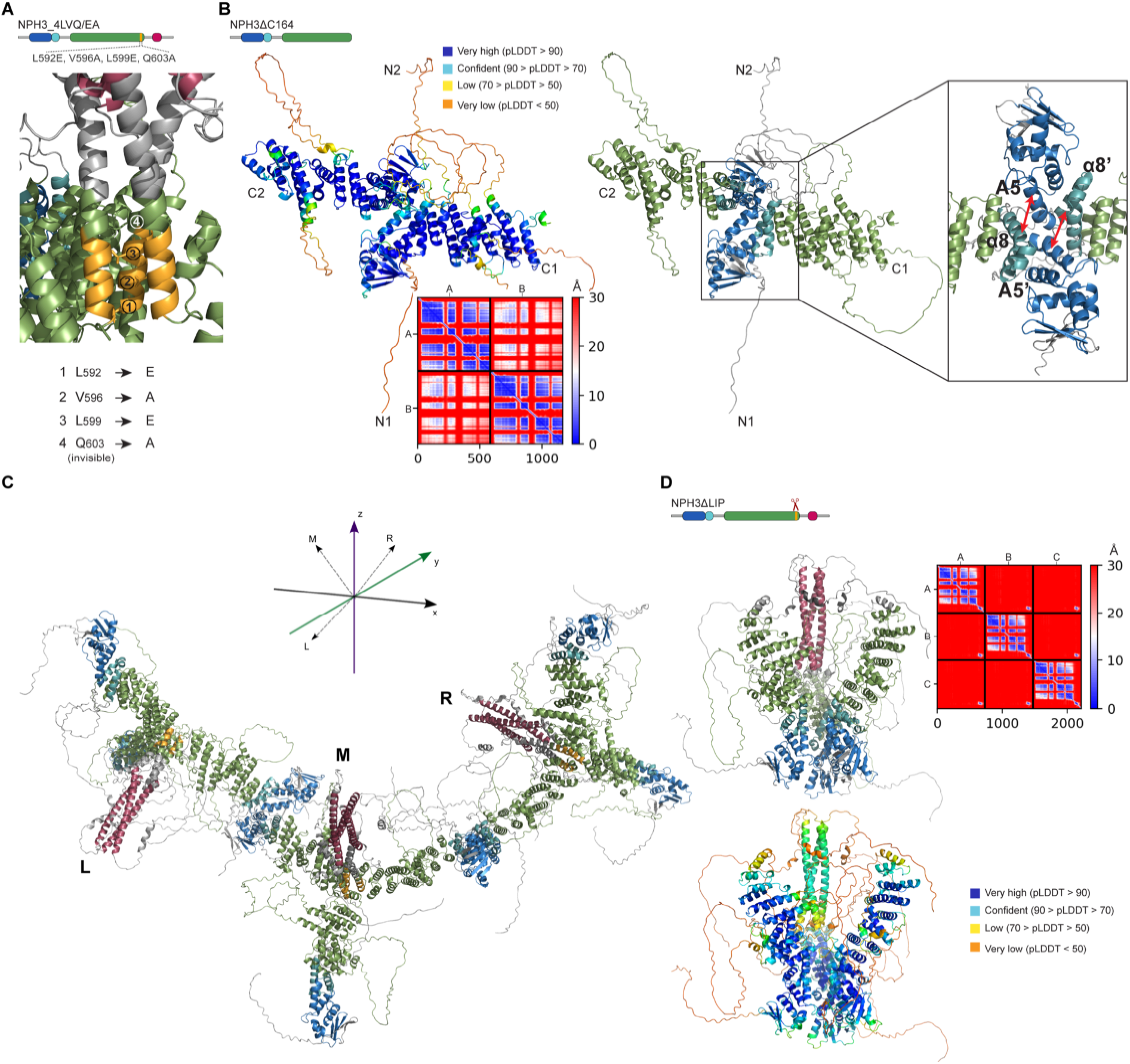
Crosslinking of NPH3 trimers via BTB-dependent antiparallel interactions could enable the formation of higher-order assemblies. (**A**) Enlarged view of the LIP helix bundle showing the position of the mutated residues. (**B**) Ribbon plot of the NPH3ΔC164 dimer predicted by AF2M and color-coded according to the pLDDT score (**left panel**) or the schematic domain representation (Fig. 1A) (**right panel**) or the pLDDT score. The corresponding PAE plot is shown in the **lower panel**. Enlarged view depicts the potential antiparallel BTB-BTB interaction based on the interaction between helix A5 of the core BTB (dark blue) and helix α8 of the C-terminal BTB extension (light blue). N1, N2, N-terminus; C1, C2, C-terminus. (**C**) Ribbon plot of three NPH3 trimers crosslinked via antiparallel BTB-BTB interaction. Note that each trimer comprises one or two additional BTB domains that could be crosslinked. The pictogram indicates the position of each CC domain in 3D space. (**D**) Ribbon plot of the NPH3ΔLIP trimer predicted by AF2M and color-coded according to the schematic domain representation (**upper panel left**) or the pLDDT score **(lower panel**). The corresponding PAE plot is shown in the **upper right panel**.

**Figure EV5.**
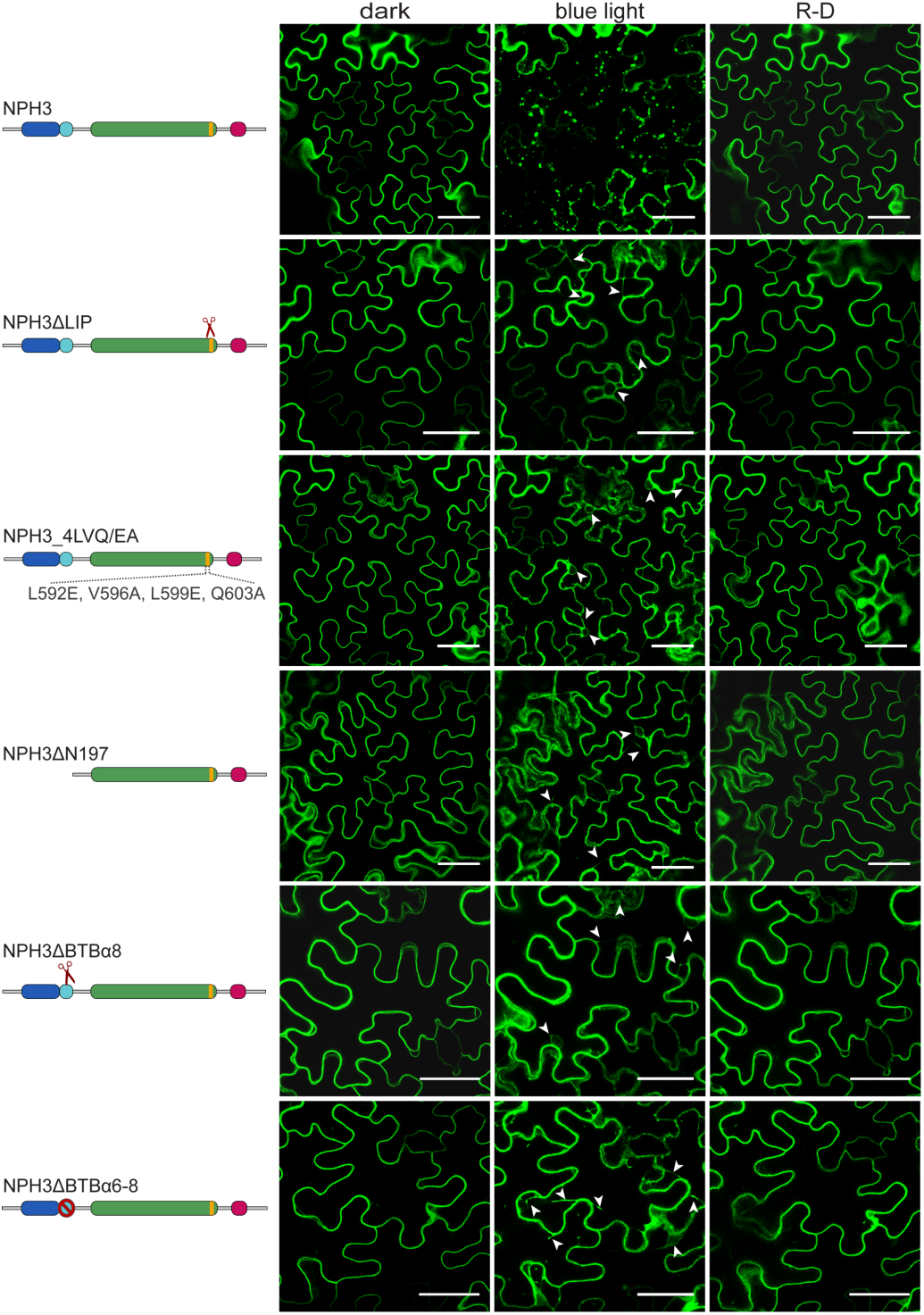
Light-dependent changes in subcellular localization are retained in condensate-incompetent NPH3 variants. Schematic representation of NPH3 variants used and representative confocal images of leaf epidermal cells from *N. benthamiana* transiently expressing (35S promoter) different GFP:NPH3 variants. Dark-adapted tobacco plants were either maintained in darkness (dark), treated with BL (∼ 11 min GFP laser) or re-transferred back to darkness (30 min) after irradiation (R-D). Scale bars, 50 µm. Experiments were repeated at least three times with similar results. White arrowheads indicate cytoplasmic strands.

**Figure EV6.**
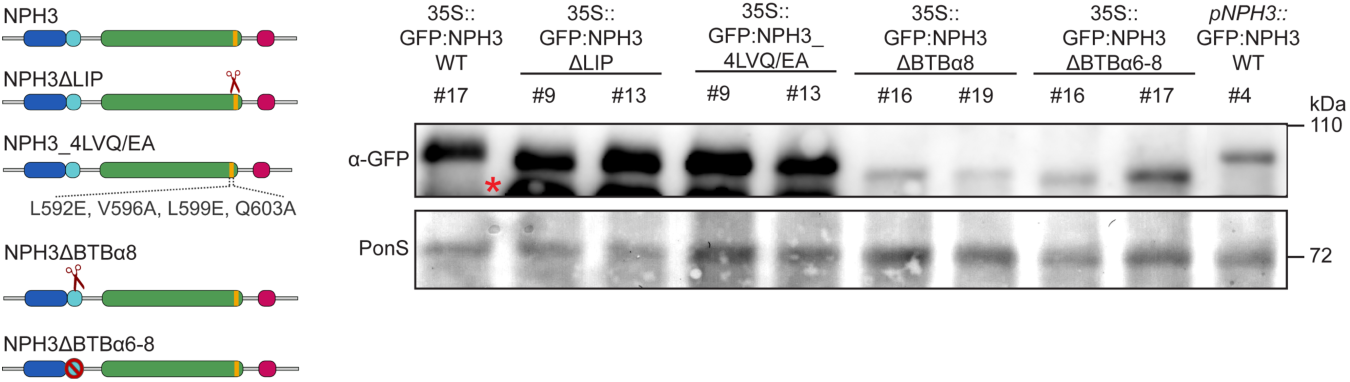
Expression level of NPH3 variants in transgenic Arabidopsis lines. Schematic representation of NPH3 variants used and immunoblot analysis of total protein extracts from 3-day-old etiolated Arabidopsis *nph3-7* seedlings expressing (35S promoter) GFP-tagged NPH3 variants. Proteins (approx. 15 µg) were separated on 8 % SDS-PAGE gels and subsequently immunodetected with an anti-GFP antibody. Red asterisk indicates degradation product. Immunodetection of native promoter driven NPH3 expression, which restores the phototropic response of *nph3-7* (Reuter *et al*, 2021; Sullivan *et al*, 2021), is shown for comparison. PonS - Ponceau Staining.

**Table EV1. Blue-light dependent NPH3 interactors in etiolated Arabidopsis seedlings**. Analysis of GFP:NPH3 immunoprecipitates via mass spectrometry based on two biological replicates. This table lists only NPH3 clients showing at least a twofold increase in normalized iBAQ values following BL irradiation. Protein intensities were converted to the relative abundance (%) of the bait protein and client proteins showing a value ≥ 1 % are highlighted in light red.

**Table EV2.**
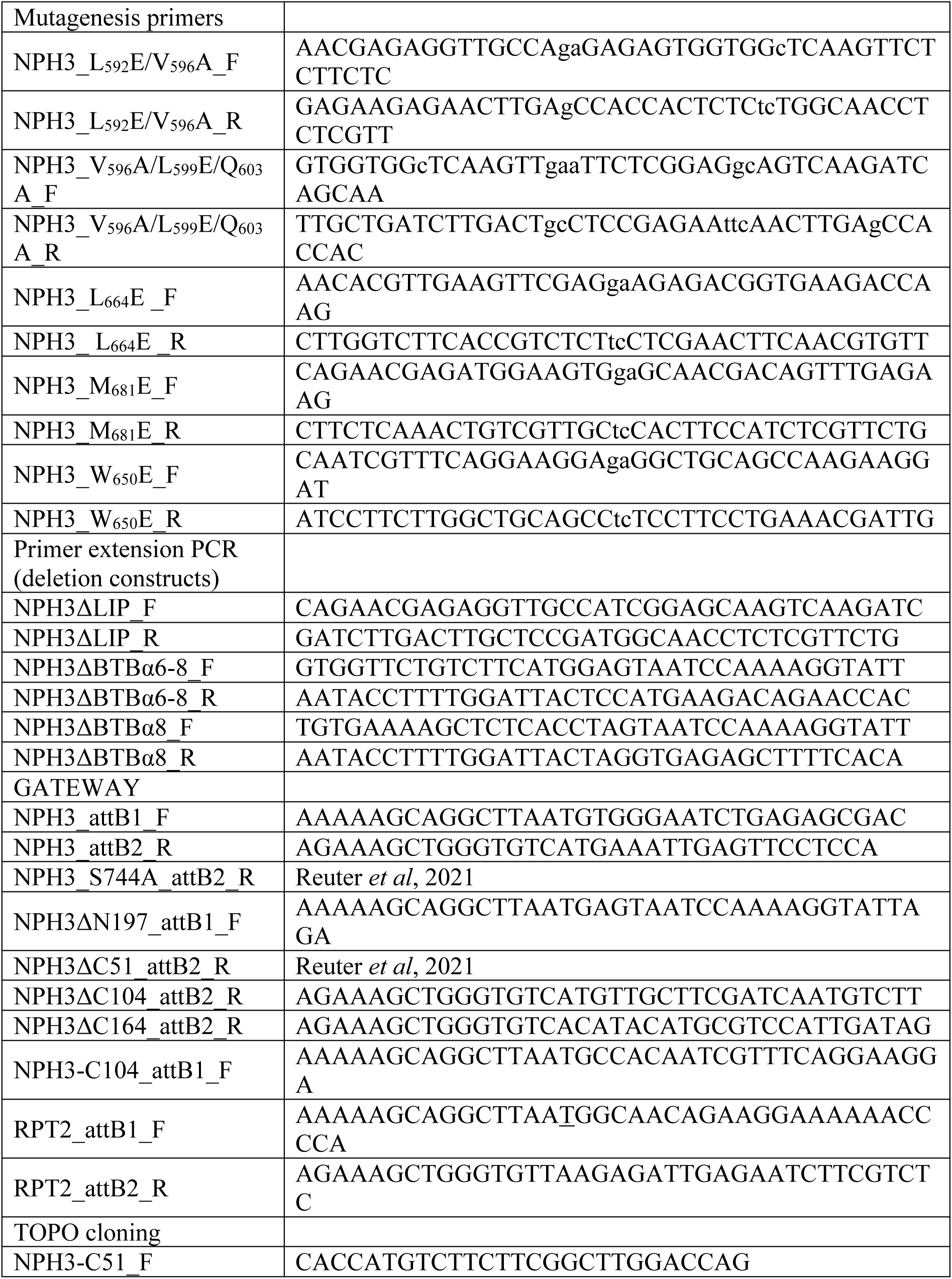
List of primers used in this study.

## Notes

### Competing Interest Statement

The authors have declared no competing interest.

### Summary of Updates

This version of the manuscript has been revised to include additional data, update several figures and sections.

